# Cell State Transitions Drive Transcriptome-Proteome Alignment in a Minimal Photoautotroph

**DOI:** 10.64898/2026.10.20.760503

**Authors:** Jordan Rozum, Xiaolu Li, Young-Mo Kim, Zachary Johnson, William Sineath, Marci R. Garcia, Natalie C. Sadler, Pavlo Bohutskyi, Margaret S. Cheung, Wei-Jun Qian, Song Feng

## Abstract

The cyanobacterium *Prochlorococcus marinus* MED4 is a minimal photoautotroph that maintains circadian regulation despite its reduced circadian regulatory toolkit relative to other cyanobacteria. Moreover, MED4, like other small prokaryotes, has low transcript counts per cell and poor transcriptome-proteome alignment. A systems-level understanding of how MED4 uses its minimal regulatory machinery to achieve a well-coordinated circadian cycle is currently lacking. We address this knowledge gap by analyzing global proteomics, transcriptomics, metabolomics, and lipidomics collected across a circadian day. Integrating these data with statistical analyses, enzyme-constrained metabolic modeling, and stochastic mathematical modeling allows us to reveal and characterize six major population-level cell states. Here we present evidence that high-light response regulator RpaB plays an important night-time regulatory role in MED4 by inhibiting RpaA, the master transcriptional activator of *dusk genes* and HimA, a regulatory target of RpaA previously shown to influence DNA conformation, is highly correlated with global transcriptome-proteome alignment (*R*^2^ = 0.67, *p <* 0.001). We construct stochastic differential equations consistent with genome-wide promotion of transcription by HimA as a plausible mechanism. Our HimA-driven model recapitulates alignment dynamics (*R*^2^ = 0.59, *p <* 0.01) as one plausible explanation. Collectively, our results elucidate how a minimal circadian regulatory program achieves robust coordination across transcriptional, translational, and metabolic levels.

## 1 Introduction

Cyanobacteria possess archetypal prokaryotic circadian regulatory systems [1, 2]. Among these, that of *Prochlorococcus marinus* MED4 (henceforth MED4) is the most rudimentary and is the focus of this study. Lacking nuclei, cyanobacteria cannot rely on the regulated transport of transcription factors through nucleopores to control the timing of gene expression. Instead, they rely on oscillations in the post-translational state of master regulators and metabolic proteins that are ultimately entrained by changes to photosythetically available photon flux [3, 4, 5, 6]. Arguably the most important of these regulatory circuits is the Kai oscillator and its readout transcription factor RpaA [3]. In the model cyanobacterium *Synechococcus elongatus* PCC 7942, the Kai oscillator consists of KaiA, KaiB, and KaiC. During the day, KaiA binds KaiC, leading to phosphorylation and activation of KaiC, allowing it to bind the kinase SasA [7]. SasA then phosphorylates RpaA, a master transcriptional regulator that activates expression of *dusk genes* [4]. Over the course of the night, a conformational change in the KaiB protein allows it to bind to and disrupt the activity of KaiC, leading to disassembly of the KaiA:KaiC:SasA complex. Simultaneously, increasingly abundant RpaB competes with RpaA for binding to SasA [8], and the phosphatase CikA actively inhibits RpaA [7]. This system is remarkably robust, and circadian gene expression can persist for days after exposure to constant light [9]. In contrast, the minimal photoautotroph MED4 possesses a much more fragile circadian regulatory system that requires daily entrainment by light variation [9]. This is because MED4, having adapted to nutrient-scarce environments by way of genome streamlining, lacks key regulators (KaiA and CikA) that confer robustness in *S. elongatus*. Moreover, the inhibitory effect of MED4 KaiB has been shown to be greatly reduced compared to that of *S. elongatus* [3]. Thus, a strong relationship between RpaB protein abundance and expression of RpaA targets is predicted to compensate for the absence or reduced effectiveness of canonical circadian regulators—but this has not been previously examined in detail. More broadly, the biological and dynamical principles that enable robust global coordination of transcription and translation using such a limited regulatory program remain poorly understood at the systems level.

Though there has been previous work in characterizing circadian oscillations of MED4 transcription and translation [10, 11, 12], the omics technologies such as proteomics and transcriptomics have since advanced noticeably in both depth and accuracy; moreover, a deeper understanding of the attenuated regulatory capacity of MED4 requires a more integrative multi-omics approach. To this end, we measured global proteomics, transcriptomics, lipidomics, and metabolomics from the same samples using the MPLEx method [13] over a 24-hour period with 2-hour intervals. We identified discrete states between which the population of cells transitions, with characteristic functional roles. Beyond a descriptive approach, we constructed and analyzed an enzyme-constrained flux balance analysis (ecFBA) model by applying the GECKO toolkit [14] to extend an existing genome-scale metabolic model from the literature [15]. In the resulting model, metabolic fluxes are limited by enzyme availability–computed from proteomics–and efficiency (*K*_*cat*_)–obtained from literature or predicted by deep learning methods. We compute reaction flux capacity aggregated at the pathway level and extract system-level metabolic activities and functions during transitions between cell states inferred from multi-omics measurements. In addition to light-response changes at dawn and dusk, we infer abrupt redistribution of metabolic capacity at midday as amino acid synthesis pathways are inhibited and translation and ribosome biogenesis pathways are activated.

Our multi-omics integration raised the question of why the correlation between transcriptomics and proteomics is so low (also observed previously [16, 10]), and whether this correlation changes over the course of the circadian cycle. Though proteins are translated from transcript templates, regulated protein degradation and competition for translational resources (e.g., ribosomes or tRNA) may partially decouple transcriptomics and proteomics. These effects are amplified in organisms like MED4 and other prokaryotes in which even highly expressed genes may produce only a few dozen transcripts per cell [16] that decay in a matter of minutes [17]. In this regime, transcription frequency–rather than steady-state abundance–is the dominant determining factor for protein production at the single cell level [16]. This resource-efficient regulatory strategy avoids overproduction of transcripts but increases sensitivity to resource competition and stochastic diffusive effects [18]. We hypothesized that increases in global transcription rate reduce these stochastic effects, thereby increasing transcriptome-proteome alignment. To test this in MED4, we developed a stochastic differential equation (SDE) model of regulated transcription and translation and coupled it to the abundance of HimA. HimA is a master regulator of DNA conformation that promotes RNAP recruitment [19], influences circadian transcription [20], and, as shown here, is strongly correlated with transcriptome-proteome alignment.

Collectively, our results reveal rapid shifts between transcriptional, translational, and metabolic cell states over the course of a circadian day and highlight regulatory changes that anticipate changes in light conditions. By coupling transcriptomics and proteomics, we provide the first evidence in MED4 for RpaB inhibition of RpaA activity. Through the integration of data and modeling, we discovered a new mechanism by which the alignment between a cell’s transcriptome and proteome can be modulated.

## 2 Methods

### 2.1 Multi-omics Experiment

#### Culture preparation

We maintained Prochlorococcus marinus MED4 (CCMP1986) in Pro99 medium [21] at 22^*°*^C under a 12h light–12h dark diel cycle, with lights on at 06:00 (with photosynthetically active radiation of ∼ 45*µ*mol photons m^*−*2^s^*−*1^) and off at 18:00. Cultures were grown in 1L glass bottles sparged with air and entrained to the diel cycle for at least 180 days prior to experiments.

We prepared a single large culture of MED4 and distributed 60mL into two 50mL bioreactors (30mL in each bioreactor to allow air and light penetration) with vent caps per replicate 24hrs before the first sampling time point. We destructively sampled three biological replicates (60mL) at each time point, dividing the sample for transcriptomics, proteomics, metabolomics (four mass spectrometry modes), and lipidomics (two mass spectrometry modes) (Suppl figure experimental workflow). Samples were harvested every 2 hours from 06:00 (6 AM) to 04:00 (4 AM) on the next day.

#### Transcriptomics

For transcriptomics, 10mL of culture was harvested by centrifugation (6500 ×*g*, 6min, 4^*°*^C) as individual replicate. Cell pellets were snap-frozen in liquid nitrogen and stored at −80^*°*^C until RNA extraction. And 1mL of supernant (culturing media) was collected and stored at −80^*°*^C for extracellular metabolome analysis. Total RNA was isolated using the RNeasy kit combined with QiaShredder homogenization columns (QIAGEN). Briefly, cells were lysed in RLT buffer supplemented with 1% *β*-mercaptoethanol and passed through QiaShredder columns for homogenization. Following addition of 70% ethanol, samples were processed according to the manufacturer’s protocol. Each biological replicate was carried through library preparation and sequencing independently. Prior to library construction, samples underwent DNase treatment and ribosomal RNA depletion. RNA integrity was verified by the sequencing facility before proceeding with library preparation. Paired-end sequencing (2 ×150 bp) was performed on an Illumina platform by Azenta US, Inc., yielding approximately 350 million reads (∼105 GB) per sample with single indexing. Sequencing reads were mapped to the Prochlorococcus MED4 reference genome. Adapter trimming and quality filtering were performed using TrimGalore (v0.6.10) with CutAdapt (v5.0), and read quality was evaluated with FastQC (v0.12.1). Gene-level read counts were generated using Rsubread (v2.20) with default parameters. Alignment quality metrics were obtained through Samtools (v1.21) and Qualimap (v2.3), with consolidated multi-sample quality reports generated via MultiQC (v1.28).

#### Proteomics

For each replicate, cellular metabolism was quenched by mixing 50mL of culture with an equal volume of 60% methanol pre-cooled to −20^*°*^C, yielding a final concentration of 30% methanol to preserve the in situ proteome, metabolome and lipidome. Cell pellets were harvested by centrifugation and rinsed once with ice-cold PBS, snap-frozen in liquid nitrogen, and stored at −80^*°*^C. For proteomics, metabolomics and lipidomics analysis, samples were processed using MPLEx protocol which extracted proteins, metabolites and lipids from single sample and enabled integrative omics analysis as described previously [13]. Briefly, cell pellets were resuspended in ice-cold Milli-Q water, then lysed by resuspension in ice-cold 2:1 chloroform:MeOH plus intermittent vortexing and sonication. Proteins, metabolites and lipids were extracted into three different phases by centrifugation at 4^*°*^C, 10000× *g* for 10min. Protein pellets were collected and dried with N2 stream, then resuspended in 100mM ammonium bicarbonate with 8M urea (pH 8). Samples were reduced by dithiothreitol and alkylated by iodoacetamide. Then, samples were diluted by 8-fold and digested with trypsin (trypsin:protein = 1:50, w/w) and 1mM CaCl2 at 37^*°*^C for 3hrs. The digested peptides were desalted by SPE C18 cleanup procedure, then subjected to LC-MS/MS analysis. Samples were analyzed by an Orbitrap Astral Mass Spectrometer coupled with a Vanquish Neo UHPLC System (Thermo Scientific, CA). An EASY-Spray HPLC column (150mm × 150*µ*m inner diameter) was used for separation (with 14 min gradient). The data-independent acquisition (DIA) mode was used. The isolation window was 2 m/z. Precursor m/z range was 380 – 980. Raw MS data were analyzed by DIA-NN against the annotated protein database we curated as described in the previous section. Fully tryptic search was conducted. Cysteine carbamidomethylation (+57.0215 Da) was enabled as a fixed modification. Methionine oxidation (+15.9949 Da) and acetylation on N-terminal (+42.0106 Da) were included as dynamic modifications. The raw MS spectral data are processed using DIA-NN [22] and the resulted LFQ intensity quantification of protein abundance are used to calculate the iBAQ values via DIAgui [23].

#### Metabolomics and Lipidomics

Half of the metabolite fraction was dried and incubated with 20 *µ*L of methoxyamine in pyridine (30 mg/mL) for 90 minutes at 37^*°*^C with shaking. The samples were then derivatized using 80 *µ*L of N-methyl-N-(trimethylsilyl)trifluoroacetamide containing 1% trimethylchlorosilane for 1 hour at 37^*°*^C under agitation. Following derivatization, the metabolite fractions were analyzed by gas chromatography-mass spectrometry (GC-MS) as reported previously [24]. Retention time calibration and spectra deconvolution were performed using Metabolite Detector. Metabolites were identified by matching spectra against the PNNL in-house metabolomics library and NIST23/Wiley 11 GC-MS spectral databases. Corresponding culture medium samples were also analyzed by the same method after complete drying of the samples.

The other half of metabolite fractions were analyzed using liquid chromatography-electrospray ionization-tandem mass spectrometry (LC-ESI-MS/MS) using a Thermo Vanquish Flex UHPLC system interfaced with a Thermo QExactive HF-X mass spectrometer as reported previously [25]. Five microliters of the extracts were injected and separated using an ACQUITY UPLC BEH HILIC column (2.1 ×100 mm, 1.7 *µ*m particle size) with a column temperature of 50 ^*°*^C and a flow rate of 300 *µ*L/min. Samples were analyzed in both positive and negative ionization modes using HCD (higher-energy collision dissociation). Full MS scan were acquired at a resolving power of 140,000 FWHM at m/z 200 with a scanning range of m/z 80–800. The data dependent acquisition (dd-MS2) parameters were used. Data analysis was done using Thermo Compound Discoverer 3.3. Spectra were aligned using an adaptive curve with a maximum retention time (RT) shift of 0.5 min and a mass tolerance of 5 ppm. Peaks were selected based on a minimum intensity of 1E6. Compounds were annotated based on isotopic pattern, RT, MS1 and/or MS2.

Extracted and dried lipids were resuspended in 200 *µ*L of 1:9 chloroform:MeOH. Ten microliters were loaded onto a guard column (Acquity UPLC CSH C18 VanGuard Pre-column, 130 Å, 1.7 *µ*m, 2.1 mm× 5 mm, Waters). Lipid separation was performed using a reverse phase column (Acquity UPLC CSH C18 column, 130 Å, 1.7 *µ*m, 3.0 mm× 150 mm, Waters) attached to a Waters Acquity UPLC H class system (Waters Corp), maintained at 42^*°*^C. The flow rate was 250 *µ*L/min. Data acquisition was conducted on an Orbitrap Velos Elite mass spectrometer (Thermo Fisher Scientific) in both positive and negative ionization modes. Full MS scans covered the m/z range of 200–1800. In-house developed software LIQUID (Lipid Informed Quantitation and Identification) [26] was used for all lipid species identifications. This was achieved by importing LC-MS/MS raw data and examining the diagnostic ion fragments and associated chain fragments in tandem mass spectra. MZMine2 [27] was used to align and gap-fill all datasets by matching unidentified features to their corresponding identified features. Peak intensities were exported for statistical analysis after all aligned features are manually verified.

### 2.2 Data Processing

#### Normalization and Filtering

We normalized the transcriptomics data as log_2_ transcripts per million (TPM), proteomics as median-centered log_2_ iBAQ [28], and metabolomics and lipidomics as log_2_ intensity. For each type of omics (and separately for each MS mode of metabolomics and lipidomics), we computed the sample-sample correlation matrices (Figure S1). In each of these data sets, we removed outlier samples with maximum replicate correlation more than three standard deviations below the mean across all replicate groups. In figures displaying merged metabolomics data, we represent each metabolite at most once by selecting the mode for which the coefficient of variation among replicates is lowest for each metabolite.

#### Gene Annotation and Aggregation

We apply gene ontology (GO) terms from the gene ontology database [29, 30] and KEGG pathways [31] to aggregate genes and their products according to function. Due to the relatively sparse functional annotation of the MED4 genome, we manually aggregated GO terms into meta-terms for enrichment analyses (Supplementary Table S1). The proteome allocation was assigned using GO terms and manually selected proteins (see partitioning.py in the Supplementary Code for implementation). Mass fraction in each proteome partition was calculated from iBAQ, assuming that iBAQ intensities are approximately proportional to abundance and computing approximate protein mass from the gene length.

#### Clustering and Enrichment

For each gene, we constructed a time-series for log_2_ TPM and log_2_ iBAQ and computed a z-score for each time point across the time series (independently for each transcript and each protein). We concatenated the time series for each gene’s transcript and protein, discarding genes that were not captured in both transcriptomics and proteomics experiments. We then clustered the resulting vectors using hierarchical clustering (see Supplementary Code for parameters used and parameter sensitivity plots). We performed enrichment analysis on each cluster using aggregated ontology terms (see Supplementary Table S1) via the hypergeometric test implemented in the Enrichr interface of GSEApy [32].

### 2.3 Mathematical Modeling

#### Enzyme-Constrained Flux Balance Analysis

We used the iSO595 [15] genome-scale metabolic model of MED4 as a baseline. We constructed enzyme-constrained flux balance analysis (ecFBA) models for each time point using the GECKO toolbox [14]. We extracted enzyme turnover numbers from the BRENDA database [33] and inferred unknown catalytic parameters using the DLKcat deep learning method [34]. After prediction from DLKcat, we then further fine-tuned these parameters according to the GECKO protocol [35], through which DLKcat predicted parameters from 17 reactions were adjusted. Metabolic fluxes were aggregated at the subsystem level by computing the sum of maximum absolute fluxes across all reactions in each subsystem. Flux distributions and maximum absolute fluxes were calculated using the Python package cobrapy [36].

#### Stochastic Transcription-Translation Modeling

We model regulated transcription and translation using Itô stochastic differential equations of the form

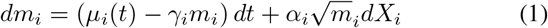

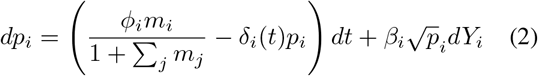

where *m*_*i*_ quantifies the abundance of the *i*^*th*^ transcript, which is translated to produce the protein of abundance *p*_*i*_. Units are arbitrary, but were selected such that *m*_*i*_ and *p*_*i*_ are on the order of molecule counts per cell [16]. Here, *X*_*i*_ and *Y*_*i*_ are vectors of independent Wiener increments so that *α*_*i*_ and *β*_*i*_ encode the strength of the noise terms. Gene activity is encoded in 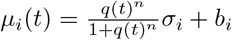 where *σ*_i_ is a randomly selected regulator promoter strength (uniform distribution) and *b*_*i*_ is a randomly selected basal transcription rate (uniform distribution). The regulatory driving force *q*(*t*) is interpolated from the iBAQ values of an input protein (HimA, in our application) scaled to the mean value across time points. Transcript decay *γ*_*i*_ is fixed at 14*h*^*−*1^ based upon an approximately 3-minute typical transcript half-life reported in [17]. Translation rate was modeled as proportional to 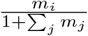 to capture competition for ribosomes. The proportionality constant *ϕ*_*i*_ was selected uniformly at random for each protein. Protein degradation is modeled as temporally regulated and varies sinusoidally with period 6*h*, 12*h*, or 24*h* (randomly selected for each protein) with uniformly random phase. For a summary of parameter values used in the simulation, see Table 1.

**Table 1:** Parameters used in simulations. Simulations include 30 batches consisting of 40 genes each. Here *U* (*S*) represents that the parameter is drawn from the uniform distribution on the set *S* independently for each value of *i* and for each batch. The parameter *δ*_0,*i*_ is similarly drawn from the normal distribution centered at 0 with standard deviation 0.2 and then clipped to the interval [−1, 1] for each *i* in each batch.

| Parameter | Value or Distribution |
| --- | --- |
| $\alpha_i$ | 5 |
| $\beta_i$ | 5 |
| $\sigma_i$ | $U([0, 1])$ |
| $b_i$ | $U([0.5, 2])$ |
| $\phi_i$ | $U([0.5, 2])$ |
| $\delta_i(t)$ | $\delta_{0,i} \sin \frac{2\pi(t-t_{i,0})}{T_i}$ |
| $\delta_{0,i}$ | $N(0, 0.2)$ clipped to $[-1, 1]$ |
| $t_{0,i}$ | $U([0, 24])$ |
| $T_i$ | $U(\{6, 12, 24\})$ |
| $\gamma_i$ | 14 |

We numerically simulated this dynamical system using the Python package sdeint [37] for *i* = 1 … 40 for 30 independent replicates. We then aggregated the results across replicates and computed the Spearman correlation between the *m*_*i*_(*t*) and *p*_*i*_(*t*) vectors at each time point. These were then averaged across each 2-hour interval centered on the input iBAQ time points. We also computed the standard error of the mean to quantify the variation in correlation over each interval.

## 3 Results

### 3.1 Multi-omics reveal major circadian and light-driven cell state transitions

We observed apparent block diagonal structure in the sample-sample correlation matrices for transcriptomics, proteomics, metabolomics, and lipidomics (see Methods), suggesting transitions between distinct cellular states. We visually grouped the blocks of samples based upon the block diagonal structure of the correlation matrices (Figure S2), and plotted the resulting groups in PCA space in Figure 2. See also Figures S3 and S4 for additional projections. These plots revealed a temporally ordered transition between grouping of samples broadly consistent with our manual aggregation based on sample-sample correlation.

**Figure 1:**
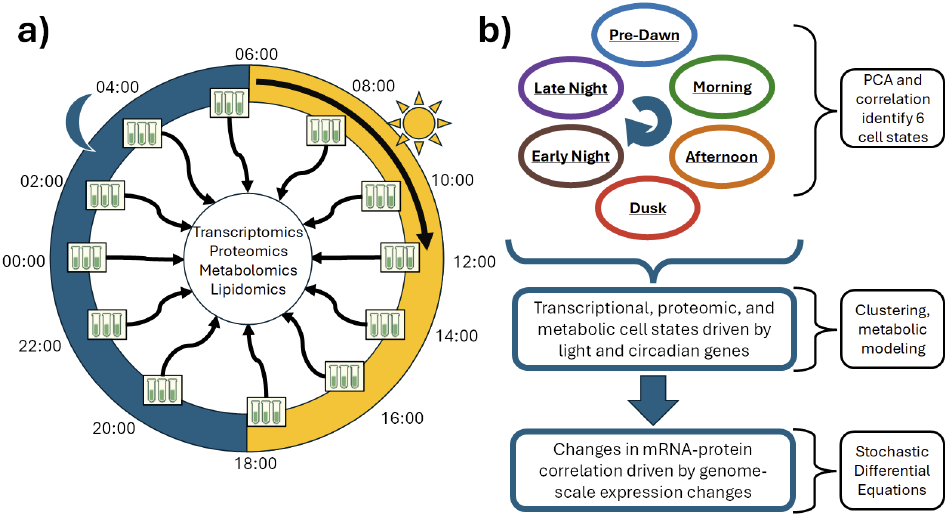
Overview of experiment design and data analysis and modeling. (a) We collected triplicate culture samples of diel-adapted MED4 every two hours over a 24-hour period, beginning at 06:00 (6 AM). Cultures were exposed to light between 06:00 and 18:00 (6 PM), and to darkness between 18:00 and 06:00 each day. We collected the 06:00 and 18:00 samples immediately before changing light conditions. We obtained transcriptomics (bulk RNA-seq), mass spectrometry proteomics, metabolomics, and lipidomics for each sample. (b) Using correlation analysis and PCA, we identified a progression of six cellular states, which we characterized using clustering and metabolic modeling. We also observed that the alignment between the transcriptome and proteome varied over time, and identified regulatory mechanisms that may drive these changes using correlative analysis and mathematical modeling.

**Figure 2:**
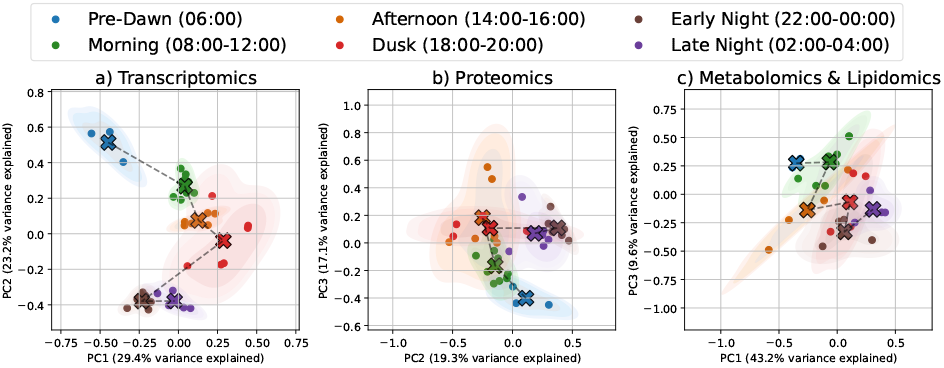
Demonstration of cell state changes in transcriptomics (a), proteomics (b), metabolomics/lipidomics (c). PCA components for projection were manually selected (see Figures S3 and S4 for alternative projections). Samples are manually grouped and colored according to the block diagonal structure preserved across sample-sample correlation matrices for each omics type (Figure S2). Samples are marked with circles, and the centroid for each group is marked with a cross. Dashed lines connect centroids for temporally adjacent groups. Shaded regions depict contours of 2-D kernel density estimates.

### 3.2 Proteomics and genome-scale metabolic modeling reveals dynamic resource reallocation

To better characterize the properties of the cell states, we used iBAQ values from bulk proteomics to estimate the time-varying allocation of proteome mass across several manually defined categories (see Methods and Supplementary Code for details). The resulting mass allocation time series are depicted in Figure 3, (see Figure S5 for a max-normalized version of this plot). We verified that in all cases, the maximum and minimum allocations are statistically significantly different (two-sample t-test, *p <* 0.05).

**Figure 3:**
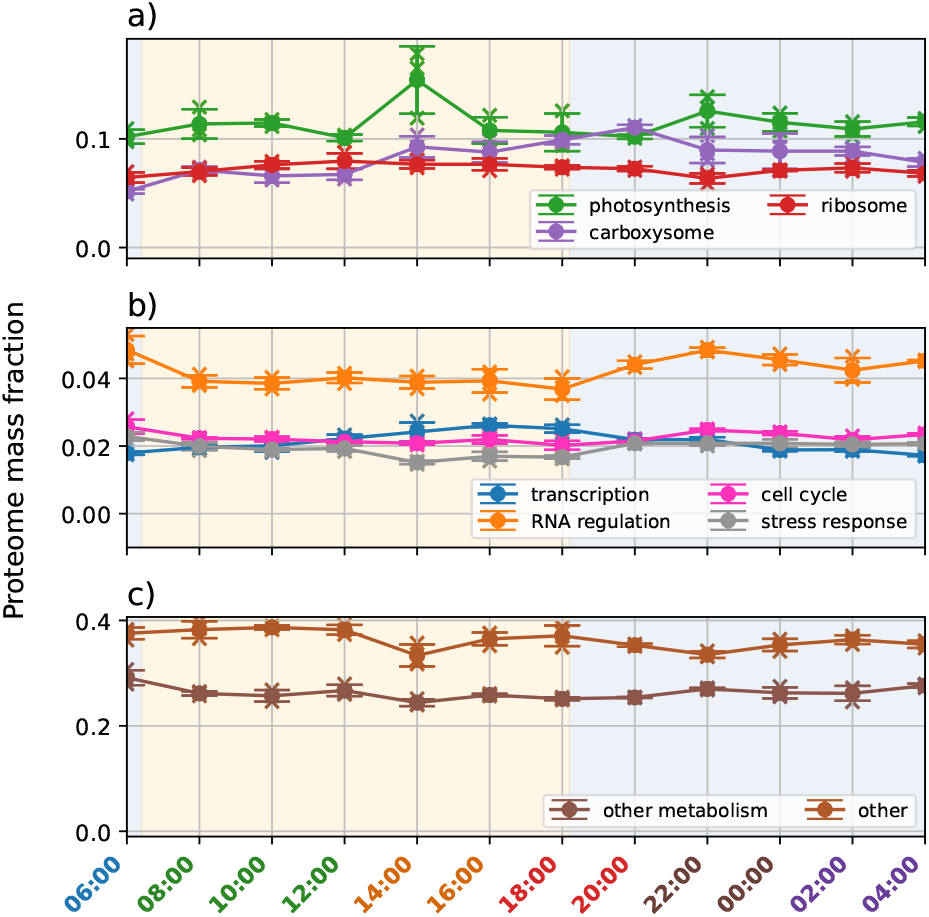
Proteome allocation shifts throughout the circadian cycle. We partitioned the proteome into nine subsets based upon functional annotation and known macromolecular complex membership. Protein mass was approximated as proportional to the product of gene length and iBAQ value, then normalized to the estimated total. Time series are plotted on separate panels (a-c) to more easily visualize individual trajectories. Time labels are colored according to manual cell state groupings. Panel backgrounds are shaded according to light condition: blue indicates dark condition, yellow indicates light condition. Dots and error bars indicate replicate means and standard deviations, and sample measurements are depicted with crosses.

As expected, we observed that the transition from 6:00 to 8:00 (“sunrise”) is marked by a rapid increase in mass allocation to carboxysome and ribosome subunits, and a modest increase in photosynthesis proteins. Remarkably, the proteome mass allocated to photosynthesis remains fairly constant, with the exception of a large (∼25 to 50%) transient increase at 14:00 in all three replicates. Throughout the day, we see an increase in ribosome allocation, peaking at the 12:00 time point, with a lagging transcriptional regulation allocation that peaks at the 16:00 and 18:00 time points, coincident with expression of “dusk” genes [4]. As the cell transitions from dusk to early evening (20:00-22:00), more of the proteome is allocated to cell cycle proteins, consistent with the nocturnal cell division of MED4 [38, 39].

We also performed clustering and enrichment analysis on the combined transcriptomic and proteomic profiles (see Methods). Enriched clusters are depicted in Figure 4. We observe dramatic expression shifts in these clusters that coincide with the cell state aggregation outlined in the previous section and is consistent with the proteome allocation of Figure 3. In particular, there is a marked increase in overall transcription following sunrise (6:00-8:00) and at dusk (18:00-20:00). We also see an increase in the protein expression for membrane transport (cluster C1 in Figure 4), central carbon metabolism (clusters C2 and C4), photosynthesis (clusters C5 and C13), lipid metabolism (cluster C12), and other systems at 14:00, which coincides with a more subtle shift in the transcription profiles of enriched clusters (and in the overall transcription profiles; see Figure 2 and Supplementary Figure S2).

**Figure 4:**
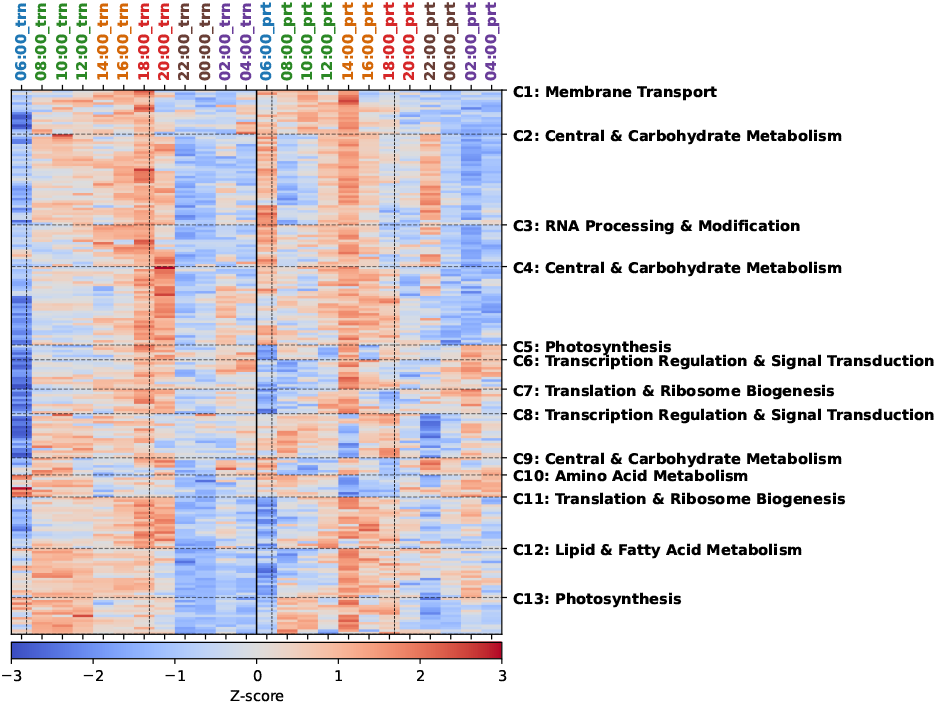
Enriched transcript-protein time series clusters. Each column represent average transcriptomic (log_2_ TPM) or proteomic (log_2_ iBAQ) quantification for a given time point, indicated by a “trn” or “prt” suffix, respectively. Time point labels are colored according to cell state groups described in the previous section. Dotted vertical lines indicate light-dark transitions; the solid vertical line separates transcriptomics columns from proteomics columns. For each omics type, rows are normalized and each cell is colored according to the resulting z-score. We grouped concatenated transcriptomic-proteomic vectors (rows) using hierarchical clustering (horizontal dotted lines) and tested for enrichment in manually aggregated GO Biological Process terms (see Methods). Only significantly enriched clusters (hypergeometric test, FDR<0.1) are depicted here. Enriched clusters are labeled C1-C13.

The metabolic fluxes of MED4 must respond to fluctuating environments and circadian-driven resource needs via a shift in transcriptional activity and downstream reallocation of proteome mass. To explore the effects of changing light and proteome allocation on metabolic flux redistribution, we turned to a genome-scale metabolic model of MED4 [15] that has been previously validated in other contexts [40]. We extended this model with enzyme constraints derived from proteomics measurements and computed maximum flux through each metabolic reaction, aggregated to the subsystem level (see Methods). Figure 5 depicts the resulting normalized subsystem-level flux capacity (rows arranged by hierarchical clustering). Subsystem capacity was defined as the sum of per-reaction maximum absolute fluxes and represents an upper bound rather than a jointly achievable flux state. Because capacities aggregate independent per-reaction maxima, Figure 5 should be interpreted as potential capacity shifts; feasibility-checked flux solutions are not shown here.

**Figure 5:**
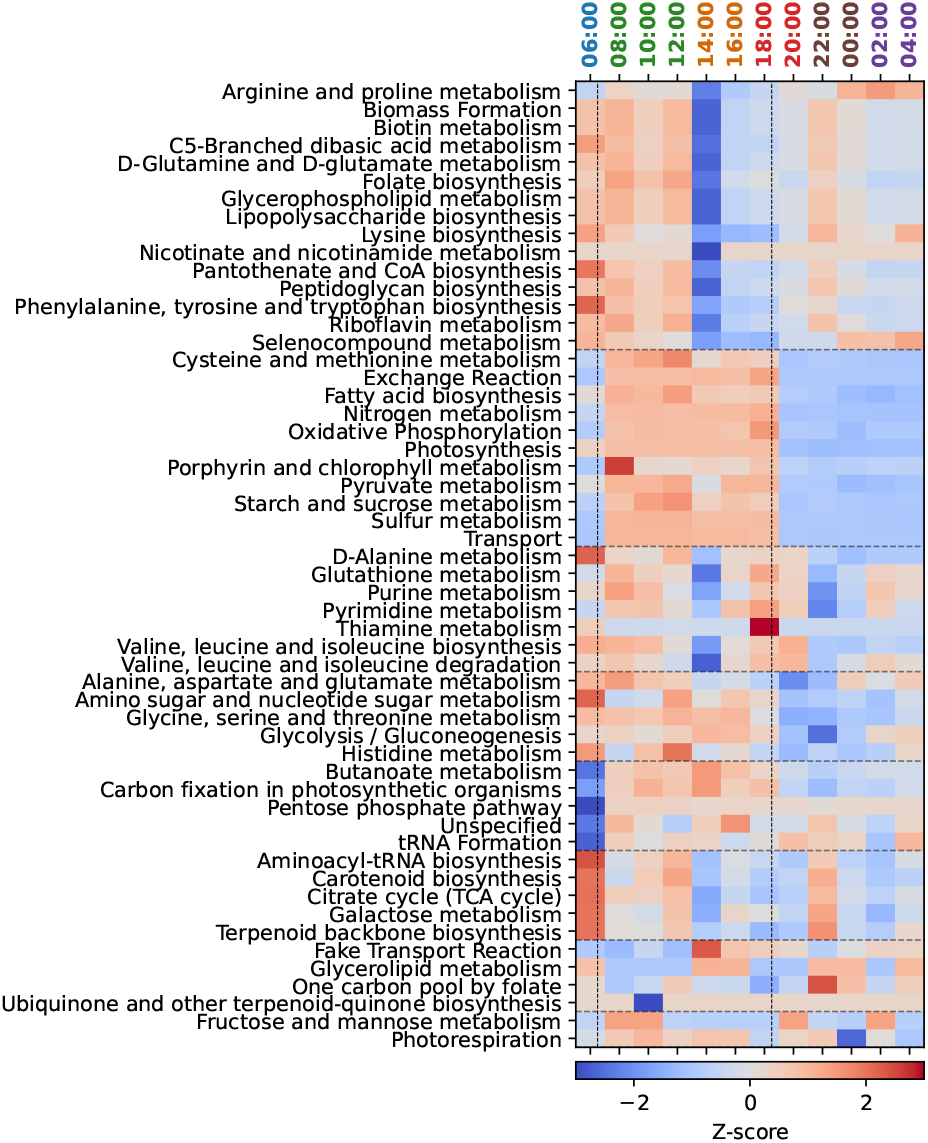
Modeled enzyme-constrained metabolic subsystem fluxes across time. Maximum absolute reaction fluxes were calculated for each reaction subject to enzyme usage constraints, stoichiometric constraints, and biomass maximization. Maximum fluxes were summed across each metabolic subsystem, then normalized across time. Time point labels are colored according to manual cell state groupings. Dotted vertical lines indicate light-dark transitions. Subsystems are organized into groups using hierarchical clustering (horizontal dotted lines), and sorted alphabetically within each cluster.

**Figure 6:**
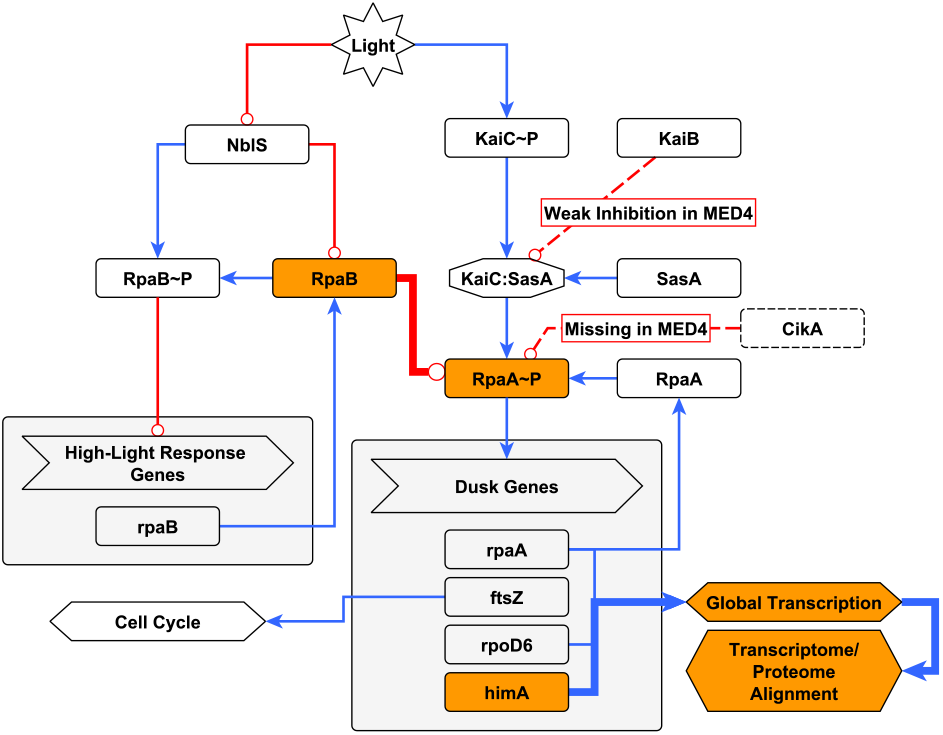
Overview of RpaA/RpaB regulatory pathway in MED4. Nodes represent biomolecular entities, protein states, processes, and external signals, distinguished by node shape. Blue arrow-tipped edges represent activating influences, and red round-tipped edges represent inhibition. Dashed lines indicate effects or entities that are absent or have reduced effectiveness in MED4 compared to other cyanobacteria. Regulatory relationships for which we present new evidence in MED4 are highlighted with orange nodes and bold edges.

Global cell state transitions (Figure 2 and time point label colors in Figure 5) coincide with major shifts in metabolic flux capacity in one or more clusters of subsystems. Modeled upper-bound changes in subsystem flux capacity (relative to the time-series mean) include: a Pre-Dawn (06:00) to Morning (08:00) increase in photosynthesis and carbon metabolism flux capacity after sunrise (approximately 1 standard deviation change); a Morning (08:00-12:00) to Afternoon (14:00-16:00) decrease in various amino acid metabolism flux capacities (approximately 3 standard deviation change); a Dusk (18:00-20:00) peak in purine and pyrimidine metabolism flux capacity (approximately 2 standard deviation change); and an Early Night (22:00-00:00) increase in amino acid metabolism flux capacities in the early portion of the night, following dusk (approximately 1 standard deviation change). The changes in amino acid and purine/pyrimidine metabolic flux capacity are offset from the day/night transition, suggesting circadian regulation of alternating metabolic functions due to limited space and resources within a small cell. Amino acid synthesis flux capacity is high just before dawn, consistent with increasing protein synthesis in preparation for heightened demand for photosystem components upon sunrise. The increased flux capacity of nucleotide precursors just before sunset is consistent with the nocturnal cell division program of MED4 [38, 39].

Many of the largest changes that are not coincident with illumination changes occur as part of the late afternoon (14:00-16:00) transitionary cell state as the cell begins to decrease photosynthetic activity prior to activation of dusk genes in the subsequent cell state. It is in this afternoon cell state that photosynthetic protein allocation is characterized by a high initial value (at 14:00) followed by a large drop (at 16:00) (Figure 3). The decrease in photosynthetic protein allocation from 14:00 to 16:00 coincides with a transitory drop in the quantification of digalactosyldiacylglycerol (DGDG) lipids (approximately 1 standard deviation decrease; see Supplementary Figure S6), which are glycerolipids present in the thylakoid and associated with photosynthetic function in cyanobacteria [41]. During this period, the metabolic flux capacity through glycerolipid metabolic pathways is increased (Figure 5), consistent with homeostatic recovery after lipid depletion.

### 3.3 HimA protein abundance coincides with transcriptome-proteome correlation, global expression, and circadian regulation

In cyanobacteria, the phosphoregulated transcription factor RpaA governs global expression of dusk genes. Its phosphorylation state is the readout of a post-translational molecular clock: the Kai oscillator. In many cyanobacteria, such as *S. elongatus*, the Kai complex has three subunits–KaiA, KaiB, and KaiC–that collectively sense changes in ATP availability. In the active state, the Kai complex binds SasA to phosphorylate RpaA, activating it [7]. The Kai-SasA complex is then disrupted by binding of conformationally altered KaiB and RpaA is actively dephosphorylated by CikA [7]. In addition, increased abundance of RpaB–in particular unphosphorylated RpaB–has been shown to inhibit RpaA phosphorylation, ostensibly by competitive binding with SasA [8]. The picture in MED4 is somewhat different. First, KaiA is not present in MED4, and KaiC activity is controlled by autophosphorylation rather than by phosphorylation by bound KaiA [3]. Second, the inhibitory effect of MED4 KaiB has been shown to be unusually weak compared to that in other cyanobacteria [3]. Finally, CikA is absent in MED4, leaving RpaB as one of the few putative inhibitors of RpaA.

We computed the number of actively expressed genes at each time point as the number of genes with log_2_ TPM values at least one standard deviation above their mean across time (Figure 7a). We observed that the peak value occurred at 18:00, coinciding with dusk and the previously reported maximum activity of the master regulator RpaA. Because the activity of RpaA is determined by its phosphorylation state, the abundance of RpaA protein is not adequate for assessing the strength of this relationship. Instead, we used the transcription levels of several previously reported [4] high-confidence RpaA regulatory targets as surrogates: *rpoD6* (sigma factor), *ftsZ* (cell division protein), *himA* (regulator of DNA conformation), and the *rpaA* gene itself (Figure 7b). The TPM time series for these four genes follow a similar overall pattern in our data set: a steady increase throughout the day, peak activity coinciding with maximal gene expression at dusk, a rapid decrease in the early night, and a recovery period in the late night. The decrease and recovery pattern in RpaA targets that we measured during the night hours suggests active inhibition of RpaA. Previous work demonstrated that RpaB abundance inhibits RpaA activity in *S. elongatus*. To assess whether a similar mechanism could be present in MED4, we plotted the RpaB abundance (log_2_ iBAQ) over time (Figure 7c). Indeed, we found increasing nocturnal RpaB abundance that coincided with decreased RpaA target transcription, and a subsequent decrease in RpaB abundance after 22:00 that coincided with the recovery of RpaA target transcription.

**Figure 7:**
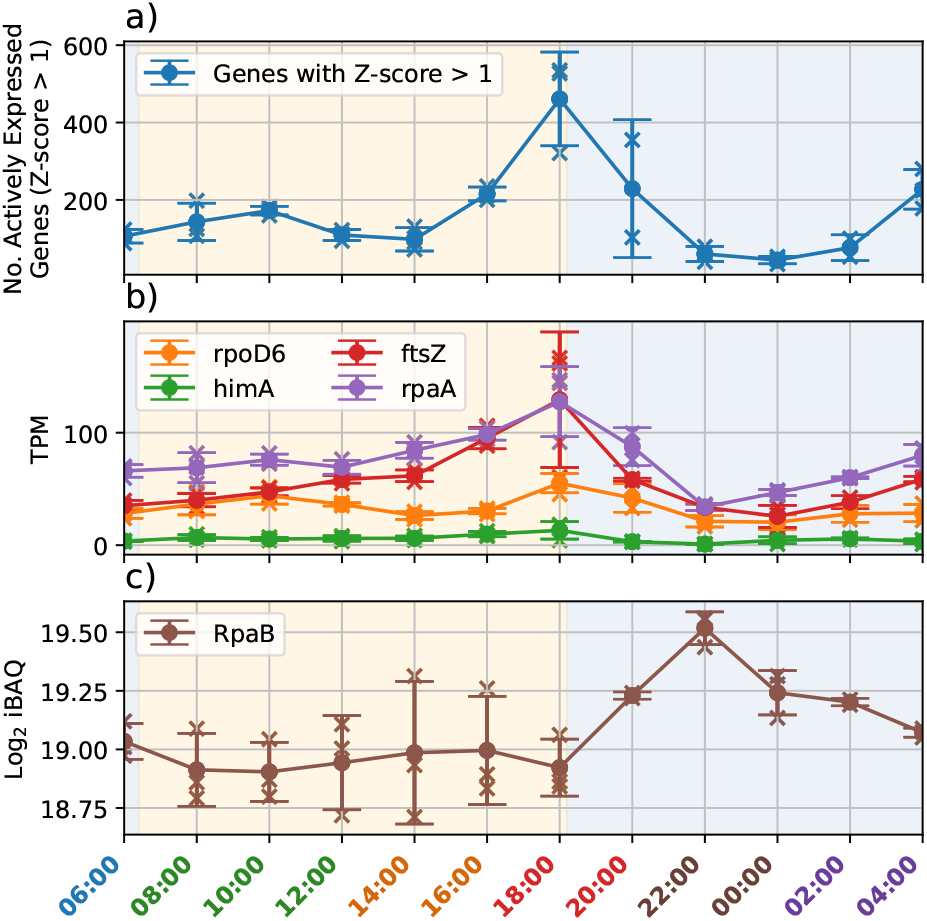
Global transcript-protein dynamics and circadian regulation. (a) Global gene expression varies throughout the circadian cycle (blue), with the most genes actively expressed at dusk. (b) The primary transcriptional regulator of dusk-expressed genes is *rpaA*, whose activity is post-translationally modified. Transcription of its regulatory targets serves as a read-out of its regulatory state. These include *rpaA* itself (purple), the cell cycle gene *ftsZ* (red), the DNA conformation regulator *himA* (green) and the sigma factor *rpoD6* (orange). (c) Protein abundance (iBAQ) of RpaB over time. RpaB protein abundance has been shown in *S. elongatus* to inhibit RpaA activity. In *S. elongatus*, several other mechanisms also contribute substantially to the down-regulation of *rpaA*, but MED4 lacks many of the corresponding genes, or the mechanisms have been shown to be weak. Time labels are colored according to manual cell state groupings. Panel backgrounds are shaded according to light condition: blue indicates dark condition, yellow indicates light condition. Dots and error bars indicate replicate means and standard deviations, and sample measurements are depicted with crosses.

In Figure 8, we quantified the strength of the observed relationships between the RpaA target transcription time series, the active expression counts, and RpaB abundance. We find statistically significant correlations in all cases (*R*^2^ ranging from 0.24 to 0.63, *p <* 0.05 or better). We note that the correlation between *himA* transcription and actively expressed gene count is much weaker than for the other three genes (*R*^2^ = 0.24 vs *R*^2^ = 0.54 to 0.63) due to outliers with low *himA* expression and actively expressed gene count; this may be due to insufficient sensitivity at low transcript counts. Overall, the correlations in Figure 8 support our qualitative observations regarding the trends in Figure 7.

**Figure 8:**
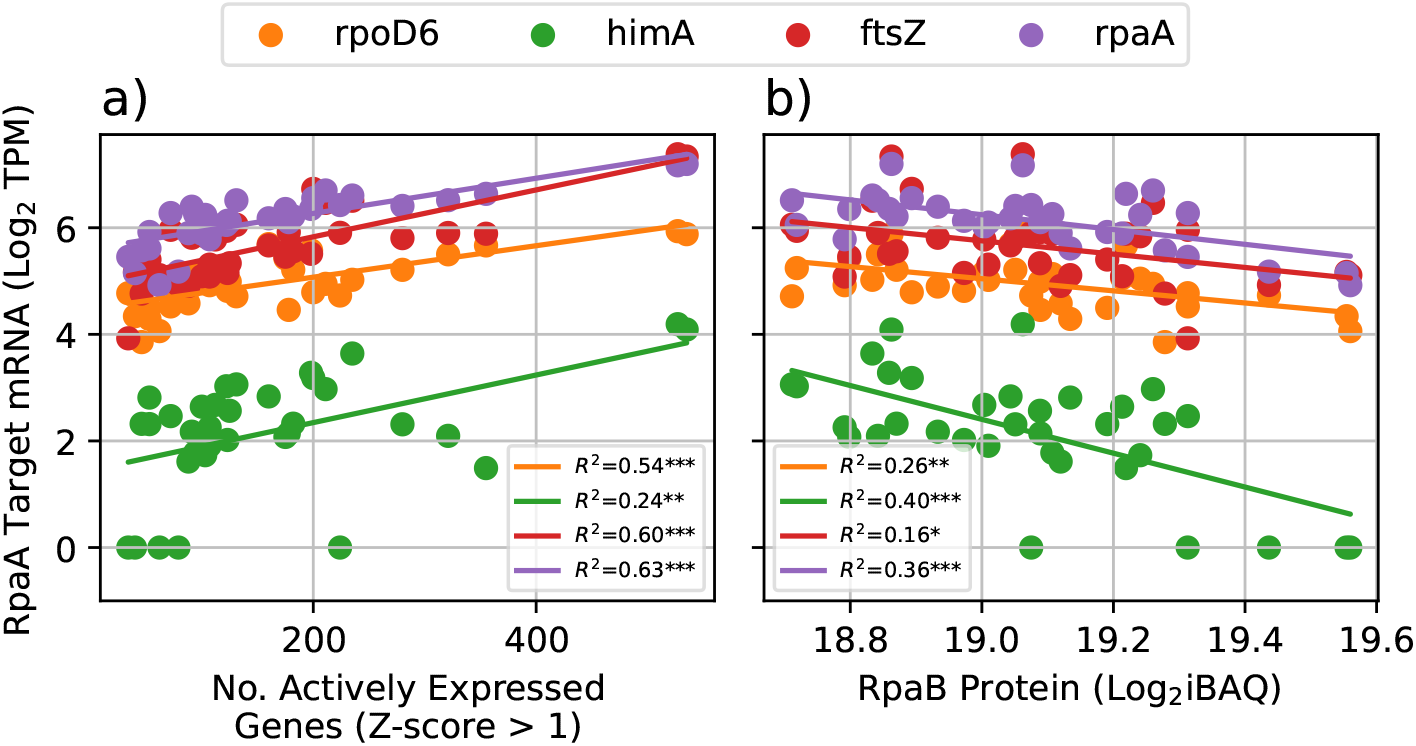
Correlation between RpaA target transcription, overall transcription (a), and RpaB abundance (b). All Pearson correlations depicted are statistically significant (*: *p <* 0.05, **: *p <* 0.01; ***: *p <* 0.001). Number of actively expressed genes (a) is calculated by normalizing log_2_ TPM across samples for each time point, independently for each gene, and counting the number of genes with z-score greater than one at each time point.

A notable feature of this and other multi-omics data sets collected in prokaryotes is an overall lack of correlation between transcript abundance and protein abundance [16, 10]. Spearman correlation of the transcript quantification with protein quantification for an individual gene can range wildly from high anti-correlation (*ρ* −*<* 0.75) to high correlation (*ρ >* 0.75), but is on average close to zero 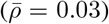 see Figure S7a for the full distribution. This correlation is not significantly increased by considering arbitrary lag times between transcription and translation (Figures S8 and S9). The Spearman correlation of a given sample’s TPM profile with its iBAQ profile–a quantity which we here refer to as *alignment*–generally falls within a much narrower range, centered on 0.10 (Figure S7b). We observed that the alignment varies temporally, peaking at dawn and dusk (Figure 9). Among proteins with known regulatory effects, we identified HimA as the most highly correlated to alignment (*R*^2^ = 0.67, *p <* 0.001; Figure 9). In other bacteria, HimA has been shown to affect global transcription and DNA conformation [20] and increase recruitment of sigma factor-bound RNAP [19].

**Figure 9:**
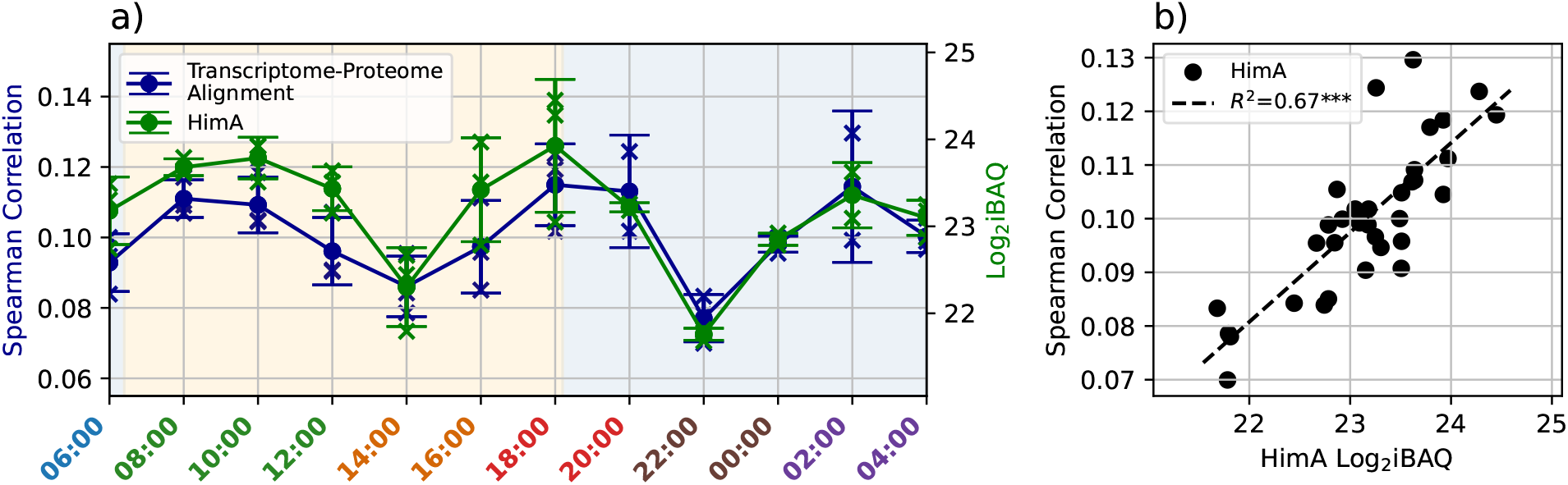
Transcriptome-proteome alignment and HimA protein abundance. Alignment (panel (a), left vertical axis) is calculated per sample as the Spearman correlation between the vector of TPM values and the vector of iBAQ values for genes captured in both transcriptomics and proteomics data sets. HimA log_2_ iBAQ (panel (a), right vertical axis) is plotted atop the alignment time series; this protein was selected as the most highly correlated to alignment. Time labels are colored according to manual cell state groupings. The background in panel (a) is shaded according to light condition: blue indicates dark condition, yellow indicates light condition. Dots and error bars indicate replicate means and standard deviations, and sample measurements are depicted with crosses. The scatter plot and regression line for transcriptome-proteome alignment vs HimA log_2_ iBAQ is depicted in panel (b) (***: *p <* 0.001).

We conjectured that global increases in transcription lead to higher signal-to-noise ratios for translation, increasing correlation between transcription and translation. To test this hypothesis in the context of HimA, we constructed a system of stochastic differential equations that model regulator-modulated transcription, ribosome-limited translation, and randomly time-varying protein degradation (see Methods). In this model, the HimA iBAQ values are used as input and influence transcription of a large number of genes. The regulatory influence of HimA does not extend to all genes in the model, nor is it fully determinative of transcription rate due to randomly selected basal transcription rates and HimA regulatory strengths. Nevertheless, as demonstrated in Figure 10, the simulated transcriptome-proteome alignment shows good agreement with the experimentally measured values (*R*^2^ = 0.59, *p <* 0.01; see Figure S10) and predicts similar mean alignment and temporal correlation as is observed experimentally (Figure S11).

**Figure 10:**
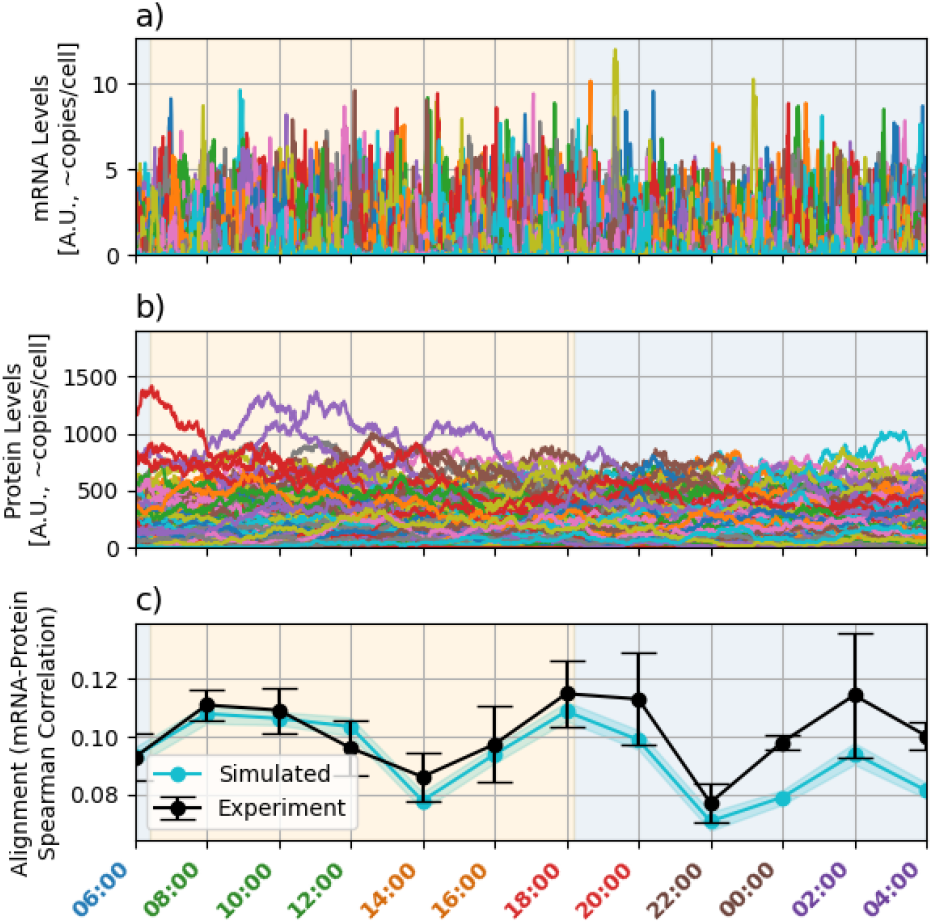
Simulation of HimA driving global transcription and transcriptome-proteome alignment. Stochastic transcription (a) is partially driven by interpolated measurements leading to downstream translation of proteins (b). Transcriptome-proteome alignment is depicted in panel (c) for both the simulation (cyan) and the experiment (black). The cyan shaded interval represents the standard error of the mean simulated alignment over the two-hour interval centered on each time point. Error bars on the experimentally measured values indicate standard deviation across biological replicates. Time labels are colored according to manual cell state groupings. Panel backgrounds are shaded according to light condition: blue indicates dark condition, yellow indicates light condition. Dots and error bars indicate replicate means and standard deviations.

## 4 Discussion

Our multi-omics measurements revealed shifts in cell state of *P. marinus* MED4, a minimal photoautotroph that temporally controls the activity of metabolic and regulatory pathways using a simple phosphorylation-driven circadian circuit. Using PCA and correlative analysis applied to transcriptomics, proteomics, metabolomics, and lipidomics data sets, we identified six population-level cell states that the system transitions between. To characterize these states, we have emphasized the redistribution of protein resources, such as metabolic enzymes, ribosomal subunits, and stress response proteins throughout a circadian day. Using enzyme-constrained genome-scale metabolic modeling, we estimated how this redistribution of proteins affects the upper-bound flux capacity of major metabolic pathways.

Collectively, our characterizations of cell state reveal not only the expected metabolic shifts in preparation for or response to changes in light conditions, but also major mid-day and mid-night shifts in metabolic allocation. For example, we see a dramatic decrease in the metabolic flux capacity of various amino acid synthesis pathways marking the transition from the morning to the afternoon cell states (between 12:00 and 14:00). This coincides with a large relative increase in the portion of the proteome allocated to photosynthetic machinery (primarily photosystems I and II). This marks the beginning of an increase in protein expression involved in translation and ribosome biogenesis. Further investigation is required to uncover the regulatory mechanisms that drive this cell state transition.

The most well-studied circadian regulatory mechanism in cyanobacteria is the Kai complex oscillator and downstream transcriptional regulation by RpaA, which is responsible for initiating cell cycle processes and the expression of dusk genes [4]. We have presented a novel multi-omics examination of RpaA activity in MED4, which possesses a simplified form of this regulatory circuit [9, 42, 2]. We show that RpaB protein abundance is anti-correlated with expression of RpaA-regulated genes, supporting the hypothesis that RpaB is the primary inhibitor of RpaA activity in MED4 (Figures 7 and 8). This mechanism has been demonstrated in *S. elongatus* [42], and is one of the few inhibitory mechanisms for RpaA that has not been shown to be diminished or absent in MED4 relative to *S. elongatus* [3].

Previous work suggested that genome streamlining as an adaptation to nutrient-poor environments has led to reduction in circadian robustness of MED4 [43] as regulatory genes are removed. In this context, the dual role of RpaB provides a resource-efficient evolutionary strategy for reducing genome size. In the daytime, RpaB acts as a high light response regulator, managing the tradeoff between photosynthetic activity and oxidative stress [8]. At night, it is co-opted to act as the primary inhibitor of RpaA, eliminating the need for dedicated inhibitor genes and thereby reducing genome size.

Generally low alignment between transcriptome and proteome is well reported in MED4 [10] and other organisms [16], owing in part to low transcript counts (on the order of a few copies per gene per cell [16]). Among proteins with known regulatory effects, we identified HimA as the most highly correlated to alignment (R2 = 0.67, p < 0.001; Figure 9), and its roles in DNA topology and RNAP recruitment motivated a parsimonious single-input model. We revealed through such mathematical modeling that an increase in overall transcription rate can explain increased transcriptome-proteome alignment, demonstrating a previously unreported principle for global regulation of the correlation between proteome and transcriptome in prokaryotes with low overall transcript counts. Using HimA protein abundance as input, we reproduced the observed transcriptome-proteome alignment in our simulations (*R*^2^ = 0.59, *p <* 0.01), supporting the mechanism specificity. Although we did not benchmark the HimA-driven model against alternative inputs (e.g., RNAP subunits, sigma factors, ribosome allocation) or negative-control surrogates, HimA’s broad regulatory effects on circadian gene expression and RNAP recruitment have been previously demonstrated, lending further support to our hypothesis [20, 19].

In this work, we have integrated transcriptomics, proteomics, metabolomics and lipidomics with mathematical and statistical modeling to improve systems-level understanding of circadian regulation in *P. marinus* MED4, a minimal photoautotroph. We identified six population-level cell states and revealed redistribution of cellular resources that is offset from dark/light transitions, indicative of circadian regulation. Our analysis of RpaA, its regulatory targets, and RpaB supports the hypothesis that RpaB, an essential high-light response regulator, moonlights as the dominant nighttime inhibitor of RpaA activity in MED4. This is consistent with genome streamlining, as MED4 lacks the CikA protein that serves a primary inhibitory role in other cyanobacteria. Most significantly, we have uncovered a new principle for global regulation of transcriptome-proteome alignment in prokaryotes whereby globally increased transcription reduces stochastic decoupling between transcript and protein abundances in organisms with low copy numbers of transcripts. We presented evidence that in MED4, the DNA conformation regulator HimA plays a major role in governing global transcription rates and, accordingly, transcriptome-proteome alignment. Together, these findings demonstrate how a minimal photoautotroph can achieve robust genome-scale coordination across its transcriptome, proteome, and metabolome.

## 5 Acknowledgments

This work is supported by the NW-BRaVE for Biopreparedness project funded by the US Department of Energy (DOE), Office of Science, Office of Biological and Environmental Research, under FWP 81832 (M.S.C.). Pacific Northwest National Laboratory is a multiprogram national laboratory operated by Battelle for the DOE under contract DE-AC05-76RL01830.

## 6 Supplementary Figures

**Figure S1:**
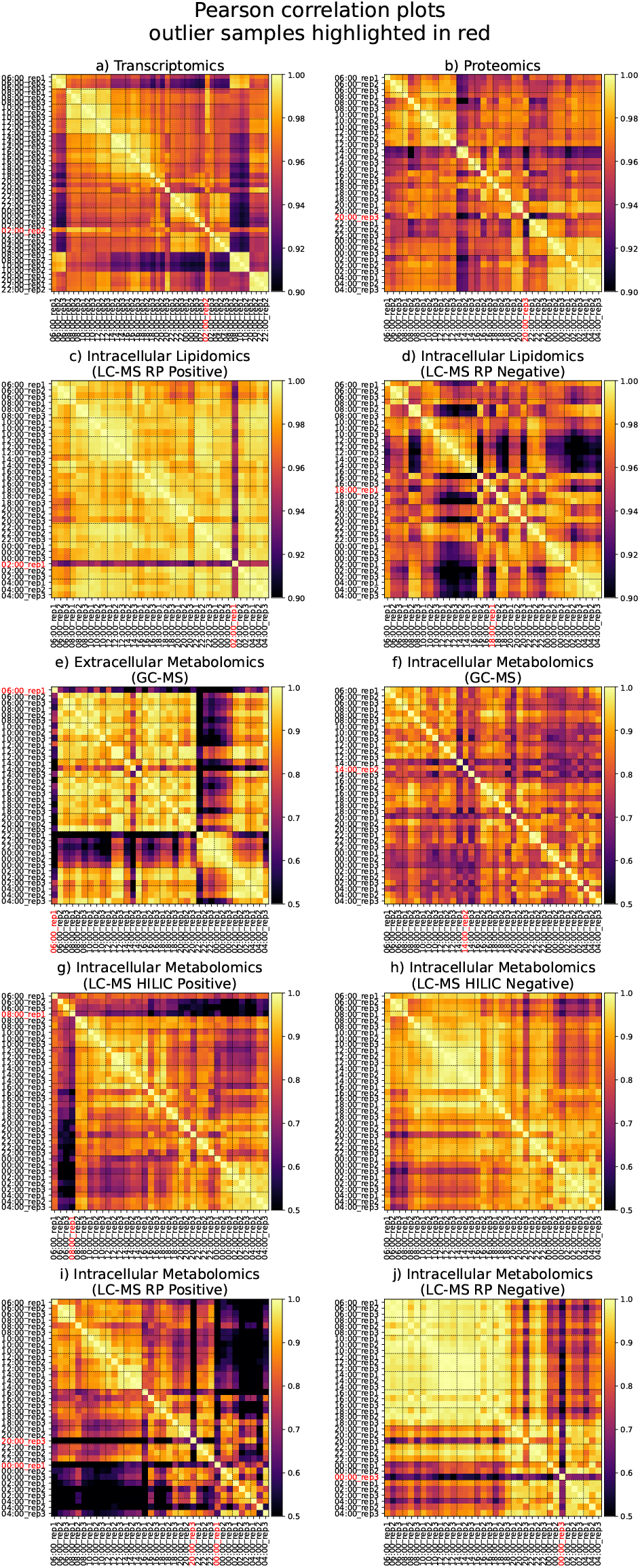
Sample-sample Pearson correlation for different omics types. Removed outlier samples in red. Black grid lines separate replicate groups.

**Figure S2:**
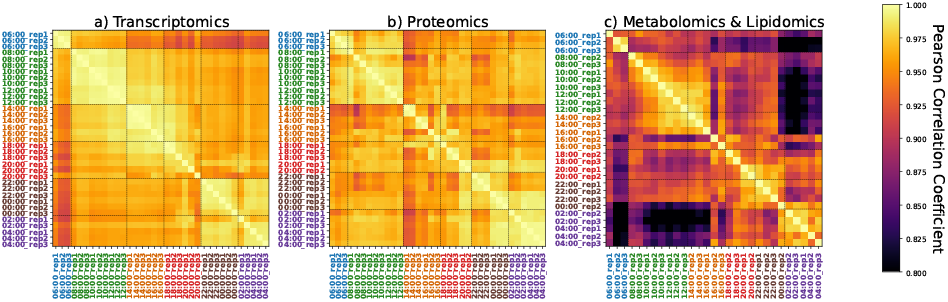
Sample-sample Pearson correlation for transcriptomics (a), proteomics (b), and metabolomics/lipidomics (c) with outliers removed. Black grid lines and label colors highlight block diagonal structure. These correspond to the same sample grouping in all three plots.

**Figure S3:**
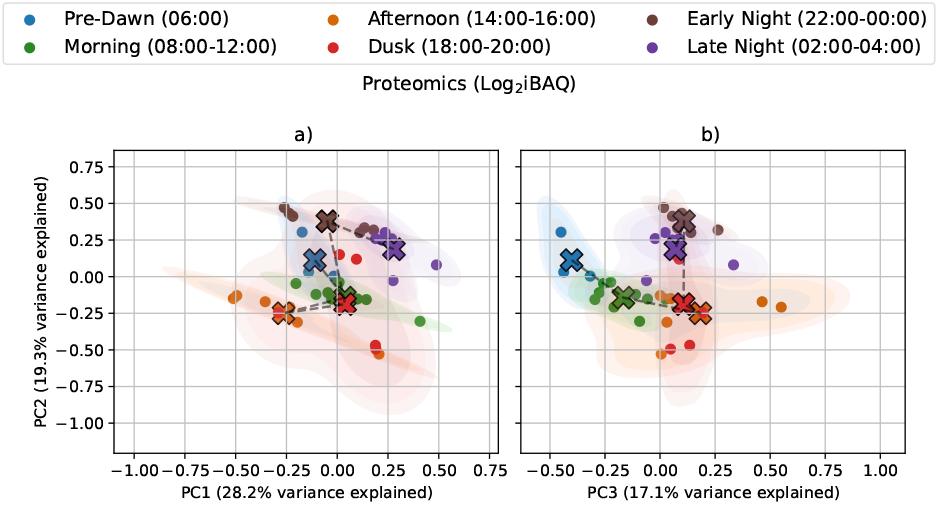
Alternative PCA projections for proteomics (log-scaled iBAQ) to complement Figure 2 in the main text. Samples are manually grouped and colored according to the block diagonal structure preserved across sample-sample correlation matrices for each omics type (Figure S2). Samples are marked with circles, and the centroid for each group is marked with a cross. Dashed lines connect centroids for temporally adjacent groups. Shaded regions depict contours of 2-D kernel density estimates.

**Figure S4:**
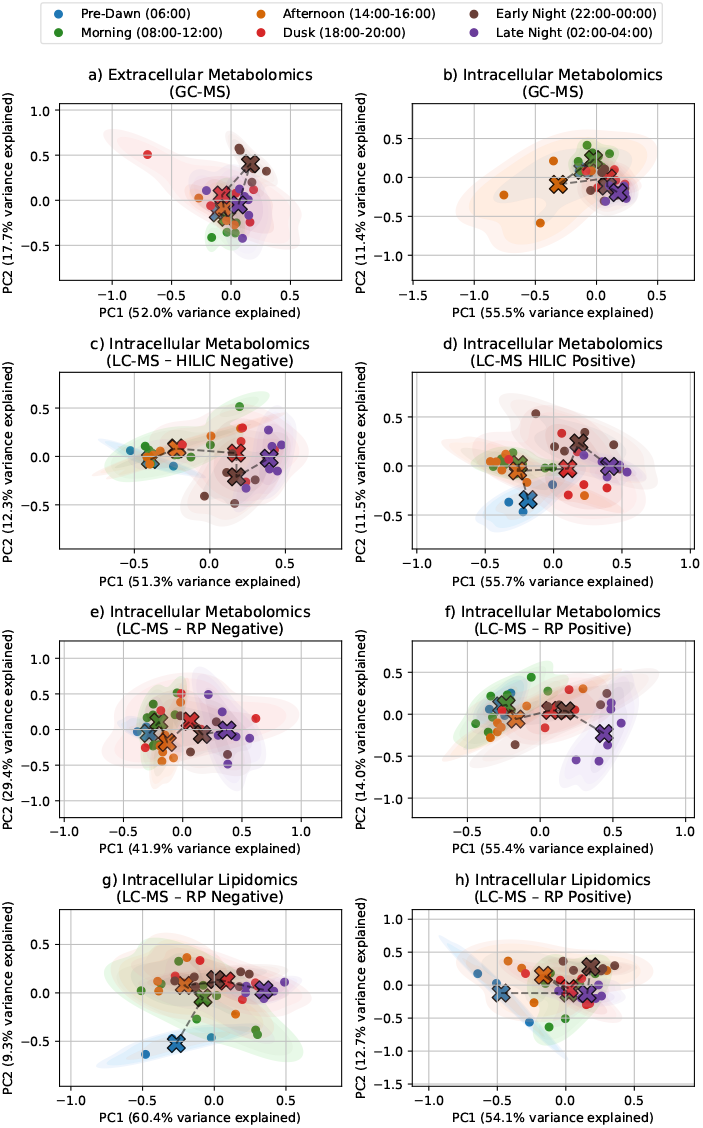
Alternative PCA projections for metabolomics and lipidomics (log-scaled), separated by technique and mass spectrometer mode, to complement Figure 2 in the main text. Samples are manually grouped and colored according to the block diagonal structure preserved across sample-sample correlation matrices for each omics type (Figure S2). Samples are marked with circles, and the centroid for each group is marked with a cross. Dashed lines connect centroids for temporally adjacent groups. Shaded regions depict contours of 2-D kernel density estimates.

**Figure S5:**
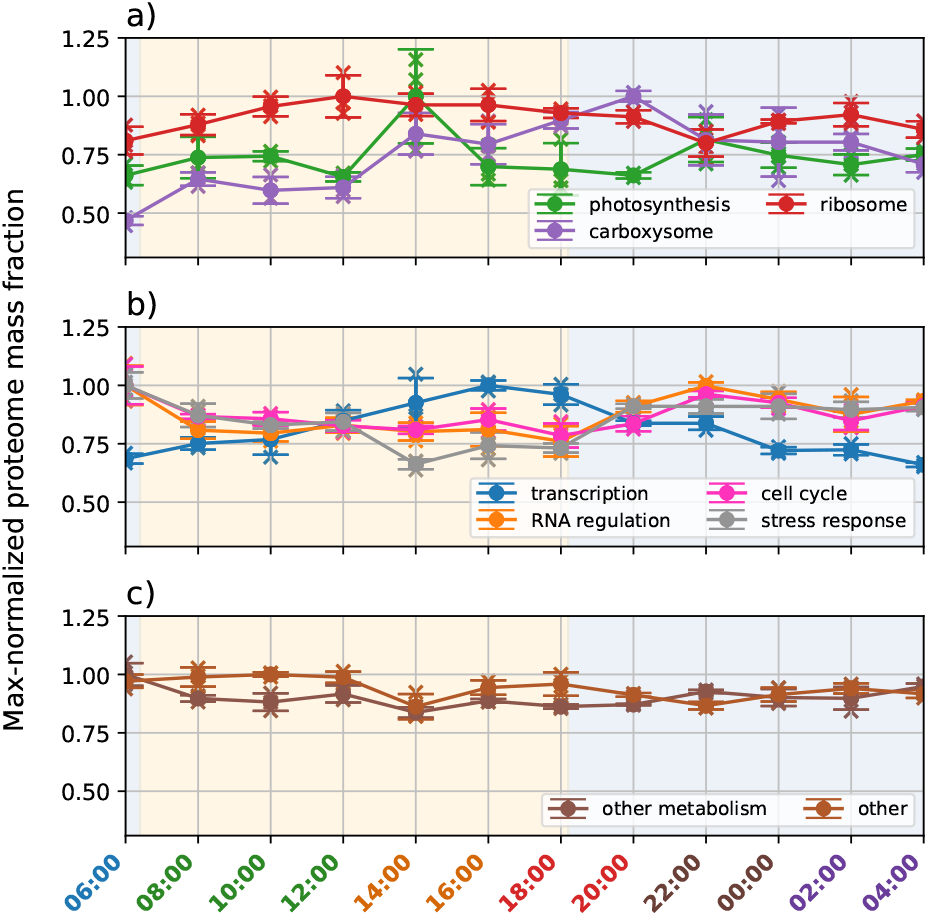
Max-normalized proteome allocation shifts throughout the circadian cycle. We partitioned the proteome into nine subsets based upon functional annotation and known macromolecular complex membership. Protein mass was approximated as proportional to the product of gene length and iBAQ value, then normalized to the estimated total. Each time series is rescaled so that the maximum mean value is equal to one. Time labels are colored according to manual cell state groupings. Panel backgrounds are shaded according to light condition: blue indicates dark condition, yellow indicates light condition. Dots and error bars indicate replicate means and standard deviations, and sample measurements are depicted with crosses.

**Figure S6:**
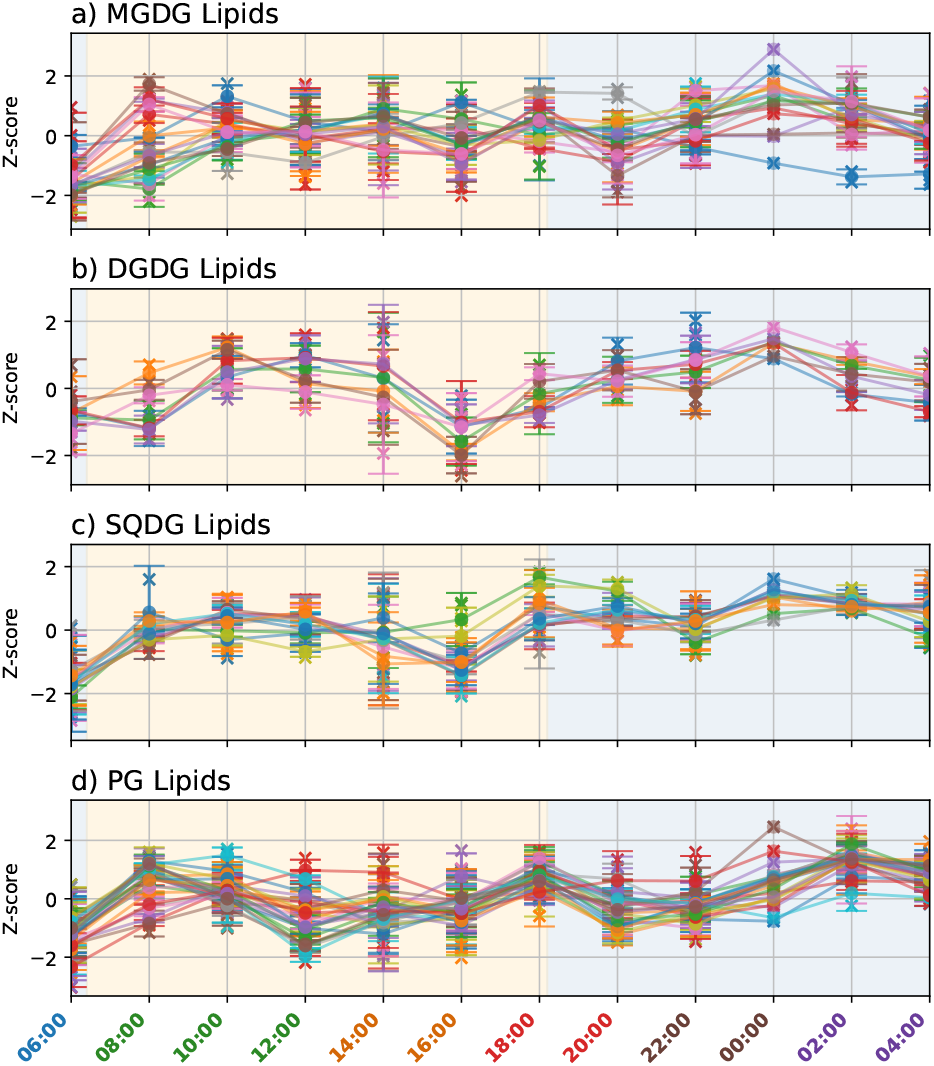
Normalized lipidomics quantification (LC-MS) time series grouped by type: a) MGDG, b) DGDG, c) SQDG, d) PG lipids.

**Figure S7:**
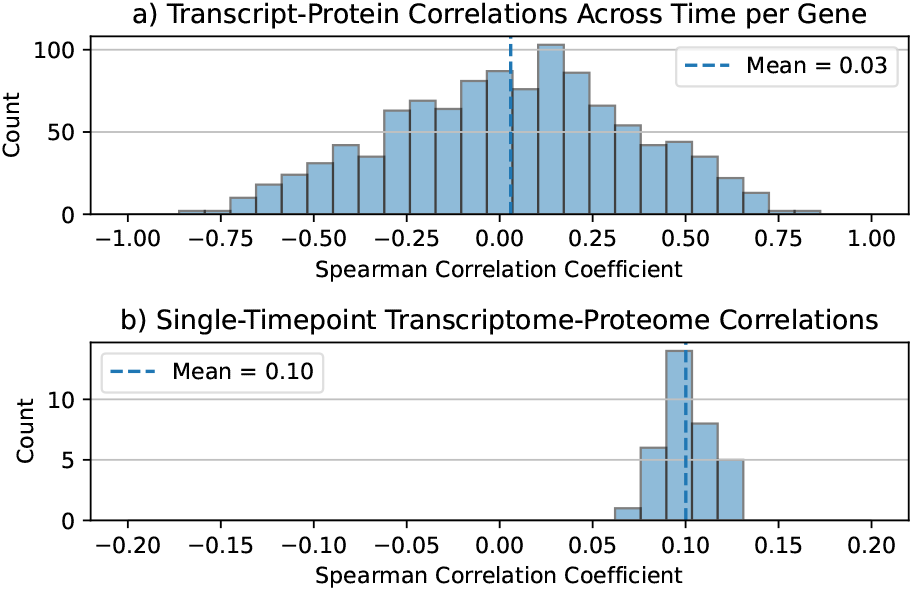
Illustration of different transcriptome-proteome correlation distributions. Average temporal correlation (a) is lower than average profile correlation (b).

**Figure S8:**
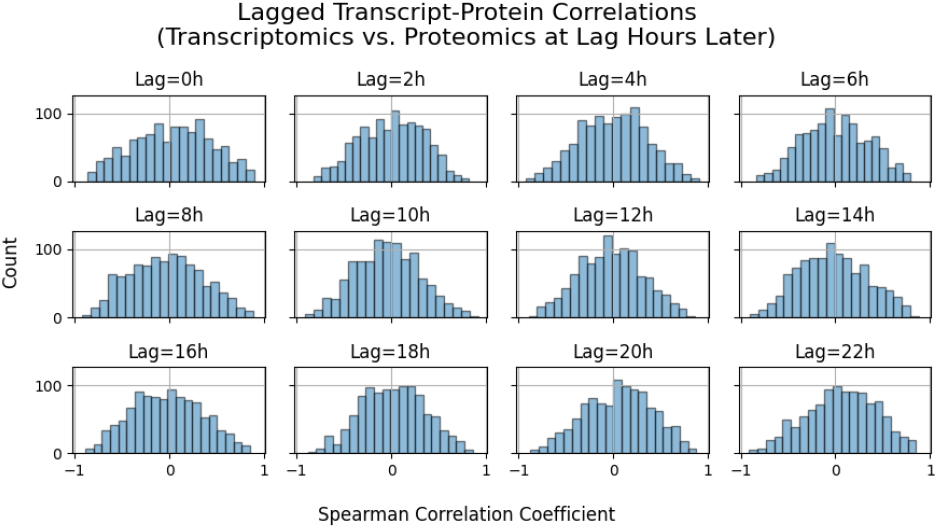
Transcriptome-proteome Spearman correlation per gene across time for different values of translational lag. We time-shifted every proteomics sample by comparing to transcriptomics samples taken at a time corresponding to two hours earlier (modulo 24 hours). Each facet corresponds to a different time shift.

**Figure S9:**
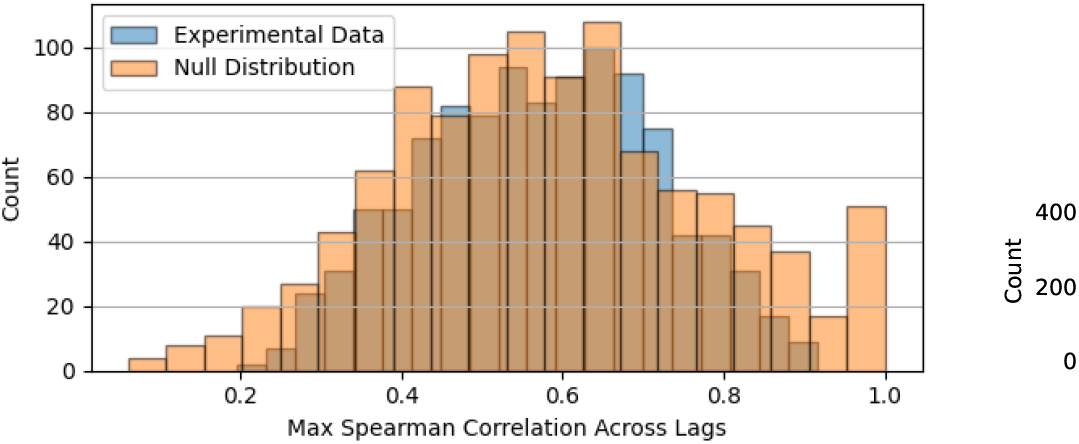
Distribution of transcriptome-proteome Spearman correlation per gene across time, maximized across lag times. The resulting distribution is compared to an equal number of samples drawn from a null distribution constructed by randomly drawing from a normal distribution with zero mean and standard deviation equal to the mean standard deviation across lagged correlation distributions. There is no statistically significant difference in the means.

**Figure S10:**
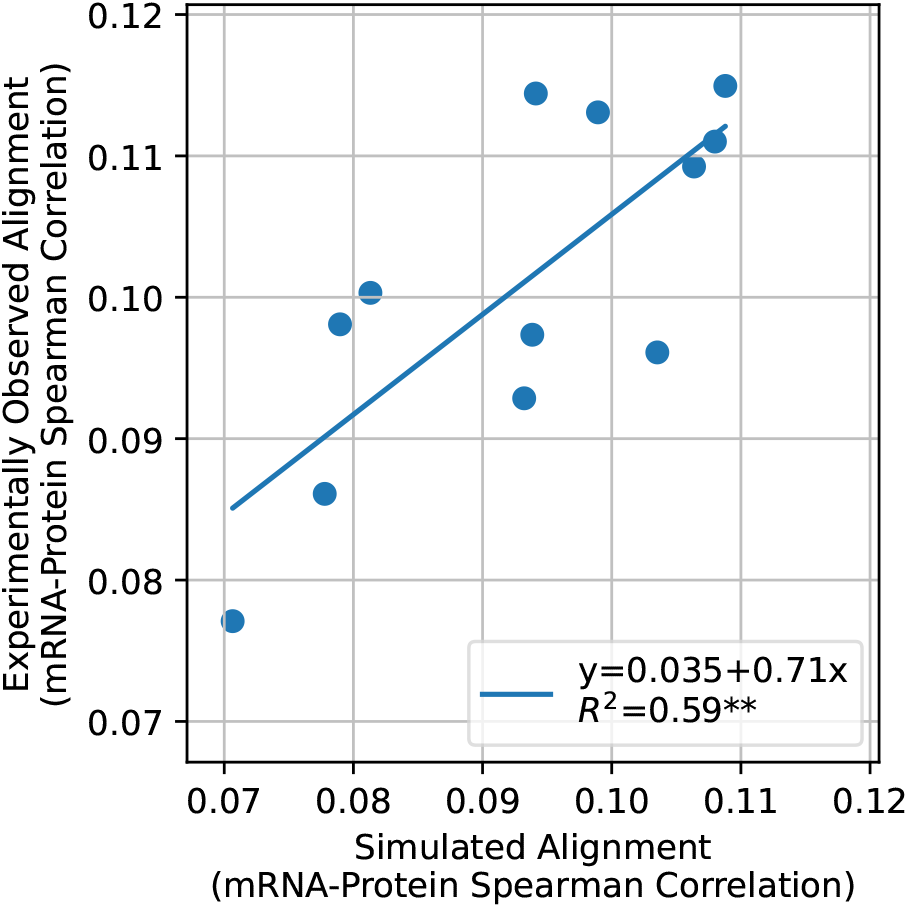
Model-experiment Pearson correlation of transcriptome-proteome alignment.

**Figure S11:**
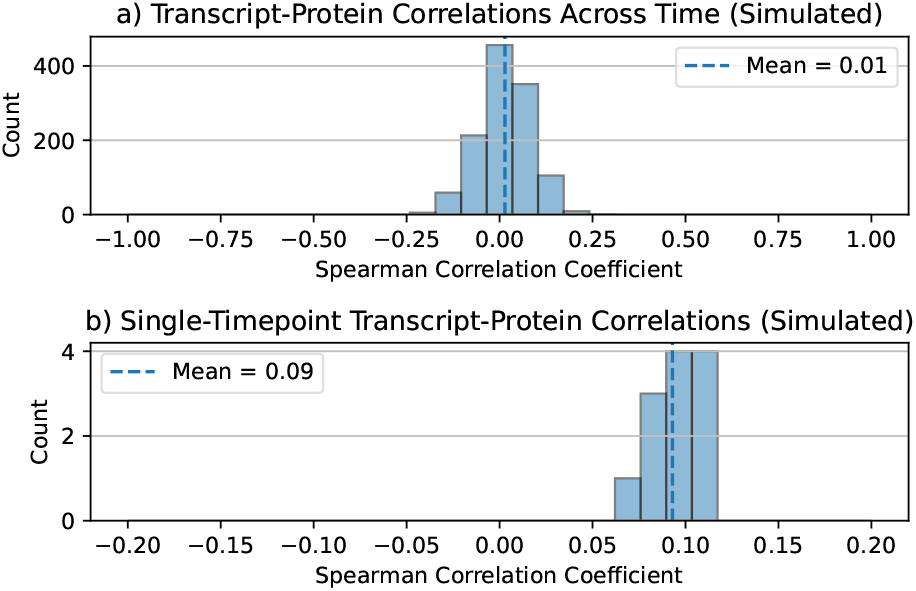
Illustration of different transcriptome-proteome correlation distributions in the simulation outputs.

## 7 Supplementary Tables

**Table S1:**
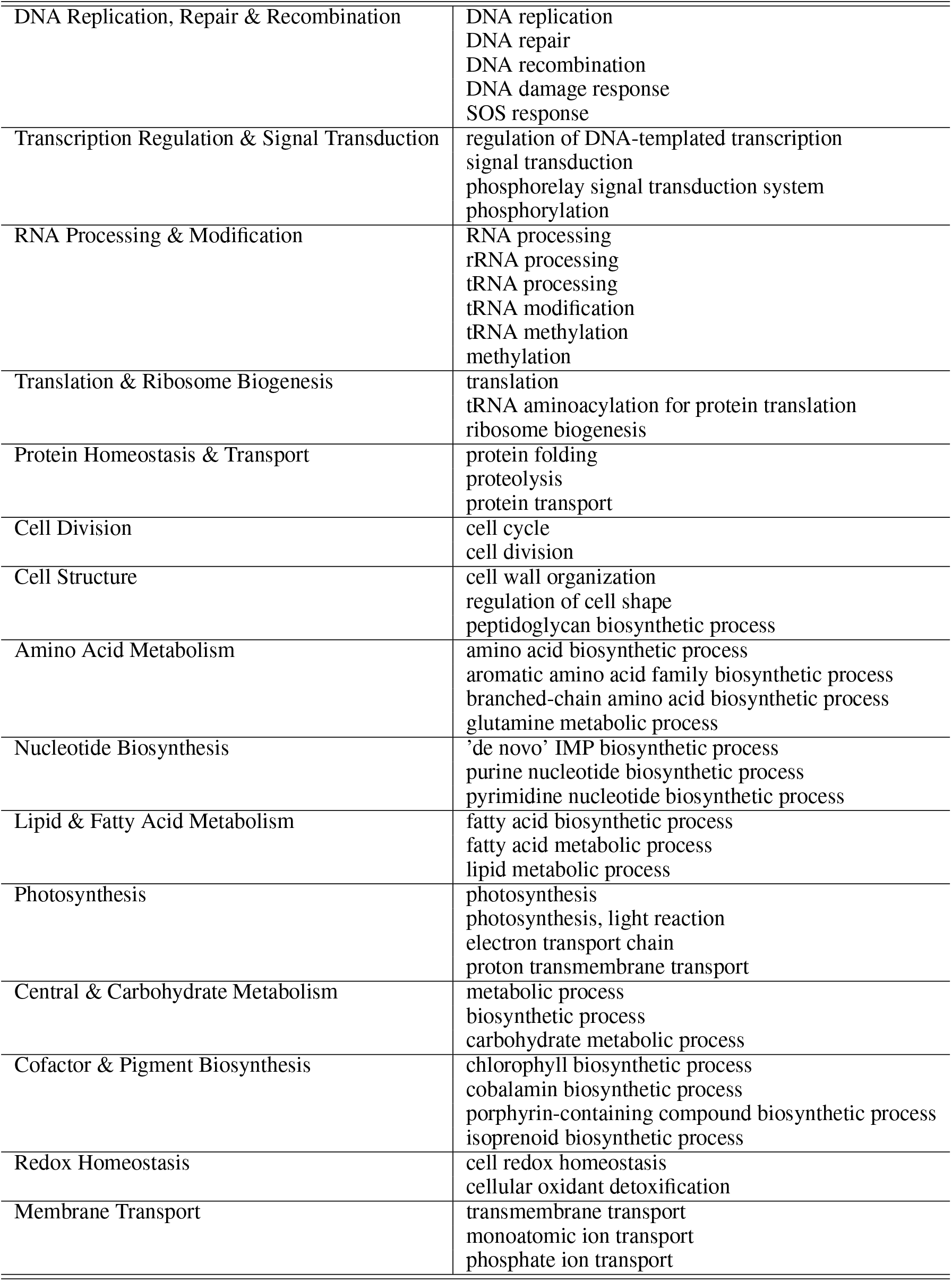
Manual aggregation of GO terms for more robust enrichment analysis of the under-annotated genes and proteins of MED4.

|  |  |
| --- | --- |
| DNA Replication, Repair & Recombination | DNA replication<br>DNA repair<br>DNA recombination<br>DNA damage response<br>SOS response |
| Transcription Regulation & Signal Transduction | regulation of DNA-templated transcription<br>signal transduction<br>phosphorelay signal transduction system<br>phosphorylation |
| RNA Processing & Modification | RNA processing<br>rRNA processing<br>tRNA processing<br>tRNA modification<br>tRNA methylation<br>methylation |
| Translation & Ribosome Biogenesis | translation<br>tRNA aminoacylation for protein translation<br>ribosome biogenesis |
| Protein Homeostasis & Transport | protein folding<br>proteolysis<br>protein transport |
| Cell Division | cell cycle<br>cell division |
| Cell Structure | cell wall organization<br>regulation of cell shape<br>peptidoglycan biosynthetic process |
| Amino Acid Metabolism | amino acid biosynthetic process<br>aromatic amino acid family biosynthetic process<br>branched-chain amino acid biosynthetic process<br>glutamine metabolic process |
| Nucleotide Biosynthesis | 'de novo' IMP biosynthetic process<br>purine nucleotide biosynthetic process<br>pyrimidine nucleotide biosynthetic process |
| Lipid & Fatty Acid Metabolism | fatty acid biosynthetic process<br>fatty acid metabolic process<br>lipid metabolic process |
| Photosynthesis | photosynthesis<br>photosynthesis, light reaction<br>electron transport chain<br>proton transmembrane transport |
| Central & Carbohydrate Metabolism | metabolic process<br>biosynthetic process<br>carbohydrate metabolic process |
| Cofactor & Pigment Biosynthesis | chlorophyll biosynthetic process<br>cobalamin biosynthetic process<br>porphyrin-containing compound biosynthetic process<br>isoprenoid biosynthetic process |
| Redox Homeostasis | cell redox homeostasis<br>cellular oxidant detoxification |
| Membrane Transport | transmembrane transport<br>monoatomic ion transport<br>phosphate ion transport |

## References

[1] Carl Hirschie Johnson, Tetsuya Mori, and Yao Xu. A Cyanobacterial Circadian Clockwork. Current biology : CB, 18(17):R816–R825, September 2008.

[2] Ilka M. Axmann, Stefanie Hertel, Anika Wiegard, Anja K. Dörrich, and Annegret Wilde. Diversity of KaiC-based timing systems in marine Cyanobacteria. Marine Genomics, 14:3–16, April 2014.

[3] Ilka M. Axmann, Ulf Dühring, Luiza Seeliger, Anne Arnold, Jens T. Vanselow, Achim Kramer, and Annegret Wilde. Biochemical Evidence for a Timing Mechanism in Prochlorococcus. Journal of Bacteriology, 191(17):5342–5347, September 2009.

[4] Joseph S. Markson, Joseph R. Piechura, Anna M. Puszynska, and Erin K. O’Shea. Circadian control of global gene expression by the cyanobacterial master regulator RpaA. Cell, 155(6):1396–1408, December 2013.

[5] Libero Gurrieri, Simona Fermani, Mirko Zaffagnini, Francesca Sparla, and Paolo Trost. Calvin–Benson cycle regulation is getting complex. Trends in Plant Science, 26(9):898–912, September 2021.

[6] Sofia Doello, Dmitry Shvarev, Marius Theune, Jakob Sauerwein, Alexander Klon, Erva Keskin, Marko Boehm, Kirstin Gutekunst, and Karl Forchhammer. Structural basis of the allosteric regulation of cyanobacterial glucose-6-phosphate dehydrogenase by the redox sensor OpcA. Proceedings of the National Academy of Sciences, 121(50):e2411604121, December 2024.

[7] Andrian Gutu and Erin K. O’Shea. Two Antagonistic Clock-Regulated Histidine Kinases Time the Activation of Circadian Gene Expression. Molecular Cell, 50(2):288–294, April 2013.

[8] Javier Espinosa, Joseph S. Boyd, Raquel Cantos, Paloma Salinas, Susan S. Golden, and Asuncion Contreras. Cross-talk and regulatory interactions between the essential response regulator RpaB and cyanobacterial circadian clock output. Proceedings of the National Academy of Sciences, 112(7):2198–2203, February 2015.

[9] Conrad W. Mullineaux and Ralf Stanewsky. The Rolex and the Hourglass: a Simplified Circadian Clock in Prochlorococcus? Journal of Bacteriology, 191(17):5333–5335, 2009.

[10] Jacob R. Waldbauer, Sébastien Rodrigue, Maureen L. Coleman, and Sallie W. Chisholm. Transcriptome and Proteome Dynamics of a Light-Dark Synchronized Bacterial Cell Cycle. PLOS ONE, 7(8):e43432, August 2012.

[11] Bang Wang, Lina Lu, Hexin Lv, Huifeng Jiang, Ge Qu, Chaoguang Tian, and Yanhe Ma. The transcriptome landscape of Prochlorococcus MED4 and the factors for stabilizing the core genome. BMC Microbiology, 14(1):11, January 2014.

[12] Erik R. Zinser, Debbie Lindell, Zackary I. Johnson, Matthias E. Futschik, Claudia Steglich, Maureen L. Coleman, Matthew A. Wright, Trent Rector, Robert Steen, Nathan McNulty, Luke R. Thompson, and Sallie W. Chisholm. Choreography of the Transcriptome, Photophysiology, and Cell Cycle of a Minimal Photoautotroph, Prochlorococcus. PLOS ONE, 4(4):e5135, April 2009.

[13] Ernesto S Nakayasu, Carrie D Nicora, Amy C Sims, Kristin E Burnum-Johnson, Young-Mo Kim, Jennifer E Kyle, Melissa M Matzke, Anil K Shukla, Rosalie K Chu, Athena A Schepmoes, et al. Mplex: a robust and universal protocol for single-sample integrative proteomic, metabolomic, and lipidomic analyses. MSystems, 1(3):10–1128, 2016.

[14] Benjamín J Sánchez, Cheng Zhang, Avlant Nilsson, Petri-Jaan Lahtvee, Eduard J Kerkhoven, and Jens Nielsen. Improving the phenotype predictions of a yeast genome-scale metabolic model by incorporating enzymatic constraints. Molecular Systems Biology, 13(8):935, August 2017.

[15] Shany Ofaim, Snorre Sulheim, Eivind Almaas, Daniel Sher, and Daniel Segrè. Dynamic Allocation of Carbon Storage and Nutrient-Dependent Exudation in a Revised Genome-Scale Model of Prochlorococcus. Frontiers in Genetics, 12, February 2021.

[16] Yuichi Taniguchi, Paul J. Choi, Gene-Wei Li, Huiyi Chen, Mohan Babu, Jeremy Hearn, Andrew Emili, and X. Sunney Xie. Quantifying E. coli Proteome and Transcriptome with Single-Molecule Sensitivity in Single Cells. Science, 329(5991):533–538, July 2010.

[17] Ju-Yuan Zhang, Wolfgang R. Hess, and Cheng-Cai Zhang. “Life is short, and art is long”: RNA degradation in cyanobacteria and model bacteria. mLife, 1(1):21–39, 2022.

[18] Zane R. Thornburg, David M. Bianchi, Troy A. Brier, Benjamin R. Gilbert, Tyler M. Earnest, Marcelo C. R. Melo, Nataliya Safronova, James P. Sáenz, András T. Cook, Kim S. Wise, Clyde A. Hutchison, Hamilton O. Smith, John I. Glass, and Zaida Luthey-Schulten. Fundamental behaviors emerge from simulations of a living minimal cell. Cell, 185(2):345–360.e28, January 2022.

[19] Raffaella Macchi, Lorena Montesissa, Katsuhiko Murakami, Akira Ishihama, Víctor de Lorenzo, and Giovanni Bertoni. Recruitment of σ54-RNA Polymerase to the Pu Promoter of Pseudomonas putida through Integration Host Factor-mediated Positioning Switch of α Subunit Carboxyl-terminal Domain on an UP-like Element*. Journal of Biological Chemistry, 278(30):27695–27702, July 2003.

[20] Vikram Vijayan, Rick Zuzow, and Erin K. O’Shea. Oscillations in supercoiling drive circadian gene expression in cyanobacteria. Proceedings of the National Academy of Sciences, 106(52):22564–22568, December 2009.

[21] Lisa R. Moore, Allison Coe, Erik R. Zinser, Mak A. Saito, Matthew B. Sullivan, Debbie Lindell, Katya Frois-Moniz, John Waterbury, and Sallie W. Chisholm. Culturing the marine cyanobacterium prochlorococcus. Limnology and Oceanography: Methods, 5(10):353–362, 2007.

[22] Vadim Demichev, Christoph B. Messner, Spyros I. Vernardis, Kathryn S. Lilley, and Markus Ralser. Diann: neural networks and interference correction enable deep proteome coverage in high throughput. Nature Methods, 17(1):41–44, January 2020.

[23] Marc-Antoine Gerault, Luc Camoin, and Samuel Granjeaud. Diagui: a shiny application to process the output from dia-nn. Bioinformatics Advances, 4(1):vbae001, January 2024.

[24] Young-Mo Kim, Shane Nowack, Millie T. Olsen, Eric D. Becraft, Jason M. Wood, Vera Thiel, Isaac Klapper, Michael Kühl, James K. Fredrickson, Donald A. Bryant, David M. Ward, and Thomas O. Metz. Diel metabolomics analysis of a hot spring chlorophototrophic microbial mat leads to new hypotheses of community member metabolisms. Frontiers in Microbiology, 6, April 2015.

[25] Mary Dixon, Antisar Afkairin, Daniel Manter, Jane Godfrey, Alison Hamm, Nathalie Munoz, Kent Bloodsworth, Vimal Kumar Balasubramanian, Cassidy Buchanan, James A. Ippolito, Meagan Burnet, and Jorge Vivanco. The root exudates of wild tomato compared to a modern variety maintain elevated soluble soil phosphorous by interacting with rhizosphere microbiota. Scientific Reports, June 2026.

[26] Jennifer E Kyle, Kevin L Crowell, Cameron P Casey, Grant M Fujimoto, Sangtae Kim, Sydney E Dautel, Richard D Smith, Samuel H Payne, and Thomas O Metz. Liquid: an-open source software for identifying lipids in lc-ms/ms-based lipidomics data. Bioinformatics, 33(11):1744–1746, June 2017.

[27] Tomáš Pluskal, Sandra Castillo, Alejandro Villar-Briones, and Matej Orešič. Mzmine 2: Modular framework for processing, visualizing, and analyzing mass spectrometry-based molecular profile data. BMC Bioinformatics, 11(1):395, July 2010.

[28] Björn Schwanhäusser, Dorothea Busse, Na Li, Gunnar Dittmar, Johannes Schuchhardt, Jana Wolf, Wei Chen, and Matthias Selbach. Global quantification of mammalian gene expression control. Nature, 473(7347):337–342, May 2011.

[29] Michael Ashburner, Catherine A. Ball, Judith A. Blake, David Botstein, Heather Butler, J. Michael Cherry, Allan P. Davis, Kara Dolinski, Selina S. Dwight, Janan T. Eppig, Midori A. Harris, David P. Hill, Laurie Issel-Tarver, Andrew Kasarskis, Suzanna Lewis, John C. Matese, Joel E. Richardson, Martin Ringwald, Gerald M. Rubin, and Gavin Sherlock. Gene ontology: tool for the unification of biology. Nature Genetics, 25(1):25–29, May 2000.

[30] The Gene Ontology Consortium. The gene ontology knowledgebase in 2026. Nucleic Acids Research, 54(D1):D1779–D1792, January 2026.

[31] Minoru Kanehisa and Yoko Sato. Kegg syntax for comparison of organisms, organism groups, and viruses by conserved gene repertoires. Protein Science, 35(6):e70652, 2026.

[32] Zhuoqing Fang, Xinyuan Liu, and Gary Peltz. Gseapy: a comprehensive package for performing gene set enrichment analysis in python. Bioinformatics, 39(1):btac757, January 2023.

[33] Julia Hauenstein, Lisa Jeske, Antje Jäde, Mathias Krull, Katrin Dümmer, Julia Koblitz, Anja Tietz, Dieter Jahn, Lorenz Christian Reimer, and Boyke Bunk. Brenda in 2026: a global core biodata resource for functional enzyme and metabolic data within the dsmz digital diversity. Nucleic Acids Research, 54(D1):D527–D534, January 2026.

[34] Feiran Li, L. Yuan, Hongzhong Lu, Gang Li, Yu Chen, Martin K. M. Engqvist, Eduard J. Kerkhoven, and Jens Nielsen. Deep learning-based kcat prediction enables improved enzyme-constrained model reconstruction. Nature Catalysis, 5(8):662–672, August 2022.

[35] Yu Chen, Johan Gustafsson, Albert Tafur Rangel, Mihail Anton, Iván Domenzain, Cheewin Kittikunapong, Feiran Li, L. Yuan, Jens Nielsen, and Eduard J Kerkhoven. Reconstruction, simulation and analysis of enzyme-constrained metabolic models using gecko toolbox 3.0. Nature protocols, 19(3):629–667, 2024.

[36] Ali Ebrahim, Joshua A. Lerman, Bernhard O. Palsson, and Daniel R. Hyduke. COBRApy: COnstraints-Based Reconstruction and Analysis for Python. BMC Systems Biology, 7(1):1–6, December 2013.

[37] Matthew Aburn and Yoav Ram. sdeint: Numerical integration of ito or stratonovich sdes. https://github.com/mattja/sdeint/tree/master/sdeint.

[38] Stéphan Jacquet, Frédéric Partensky, Dominique Marie, Raffaella Casotti, and Daniel Vaulot. Cell cycle regulation by light in<i>prochlorococcus</i> strains. Applied and Environmental Microbiology, 67(2):782–790, 2001.

[39] Riyue Liu, Yaxin Liu, Shichang Liu, Ying Wang, Kim Li, Ning Li, Daiying Xu, and Qinglu Zeng. Three-dimensional superresolution imaging of the ftsz ring during cell division of the cyanobacterium <i>prochlorococcus</i>. mBio, 8(6):10.1128/mbio.00657–17, 2017.

[40] Jordan C. Rozum, William Sineath, Pavlo Bohutskyi, Jordan Quenneville, Doo Nam Kim, Connah Johnson, Angad P. Mehta, James Evans, David Pollock, Wei-Jun Qian, Margaret S. Cheung, Ruonan Wu, and Song Feng. Synergy and antagonism in a genome-scale model of metabolic hijacking by bacteriophages. Science Advances, 12(14):eaeb7646, April 2026.

[41] Isamu Sakurai, Naoki Mizusawa, Hajime Wada, and Naoki Sato. Digalactosyldiacylglycerol is required for stabilization of the oxygen-evolving complex in photosystem ii. Plant Physiology, 145(4):1361–1370, December 2007.

[42] Mitsumasa Hanaoka, Naoki Takai, Norimune Hosokawa, Masayuki Fujiwara, Yuki Akimoto, Nami Kobori, Hideo Iwasaki, Takao Kondo, and Kan Tanaka. RpaB, Another Response Regulator Operating Circadian Clock-dependent Transcriptional Regulation in Synechococcus elongatus PCC 7942 *. Journal of Biological Chemistry, 287(31):26321– 26327, July 2012.

[43] Julia Holtzendorff, Frédéric Partensky, Daniella Mella, Jean-François Lennon, Wolfgang R. Hess, and Laurence Garczarek. Genome Streamlining Results in Loss of Robustness of the Circadian Clock in the Marine Cyanobacterium Prochlorococcus marinus PCC 9511. Journal of Biological Rhythms, 23(3):187– 199, June 2008.

